# Csf1r-mediated depletion of midbrain microglia prevents dopaminergic neuron loss during chronic colitis

**DOI:** 10.64898/2026.01.23.700559

**Authors:** Rebecca Katharina Kutscherauer, Iris Stolzer, Emely Elisa Neumaier, Mark Dedden, Pavel Kielkowski, Wei Xiang, Alexander Grotemeyer, Marco Prinz, Takahiro Masuda, Klaus-Peter Knobeloch, Veit Rothhammer, Sebastian Zundler, Johannes C.M. Schlachetzki, Jürgen Winkler, Claudia Günther, Patrick Süß

**Affiliations:** Department of Molecular Neurology, Universitätsklinikum Erlangen, Friedrich-Alexander-Universität Erlangen-Nürnberg, Erlangen, Germany; Department of Medicine 1, Universitätsklinikum Erlangen, Friedrich-Alexander-Universität Erlangen-Nürnberg, Erlangen, Germany; Department of Neurology, Universitätsklinikum Erlangen, Friedrich-Alexander-Universität Erlangen-Nürnberg, Erlangen, Germany; Department of Chemistry, Ludwig-Maximilians-Universität München, München, Germany; Institute of Neuropathology, Medical Faculty, University of Freiburg, Freiburg, Germany; Signalling Research Centres BIOSS and CIBSS, University of Freiburg, Freiburg, Germany; Division of Molecular Neuroimmunology, Medical Institute of Bioregulation, Kyushu University, Fukuoka, Japan; Department of Neurosciences, University of California, San Diego, USA; Deutsches Zentrum Immuntherapie, Universitätsklinikum Erlangen, Erlangen, Germany

**Keywords:** Microglia, colony-stimulating factor 1 receptor, gut-immune-brain axis, inflammatory bowel disease, Parkinson’s Disease

## Abstract

Inflammatory bowel disease (IBD) predisposes to neuropsychiatric comorbidity and particularly increases the risk of Parkinson’s Disease (PD) in later life. Although the gut-immune-brain axis was proposed as a link between IBD and PD and a driver of PD immunopathogenesis, the regional pattern and single-cell landscape of the brain immune response during colitis and its contribution to PD pathology remain poorly defined. Here, we observe a loss of dopaminergic neurons in the substantia nigra pars compacta of adult mice with chronic colitis. By confocal microscopy and integrated multi-omics, we reveal a complex midbrain-centered immune response to chronic colitis in comparison to the cortex, hippocampus, and striatum. Single-cell mapping of the midbrain immune landscape showed an inflammatory shift of microglial clusters including an expansion of interferon-response microglia, CD8^+^ T cell extravasation, and increased numbers of vessel-associated neutrophils. Selective myeloid cell depletion using a colony stimulating factor 1 receptor (Csf1r) inhibitor after colitis onset reduced midbrain microglia by 67% and led to a complete rescue of dopaminergic neuron loss, without affecting mucosal pathology or T cell and neutrophil migration to the midbrain. Collectively, within the complex innate and adaptive midbrain immune response to chronic colitis, we demonstrate a causal role of Csf1r-dependent microglia for dopaminergic neurodegeneration. Thus, Csf1r inhibition in IBD may not locally ameliorate colitis, but provide neuroprotection to dopaminergic neurons. These results reveal a novel cellular link between chronic gut-derived peripheral inflammation and midbrain vulnerability and thereby substantially enhance our understanding of the risk for PD related to the gut-immune-brain axis.

Graphical abstract.

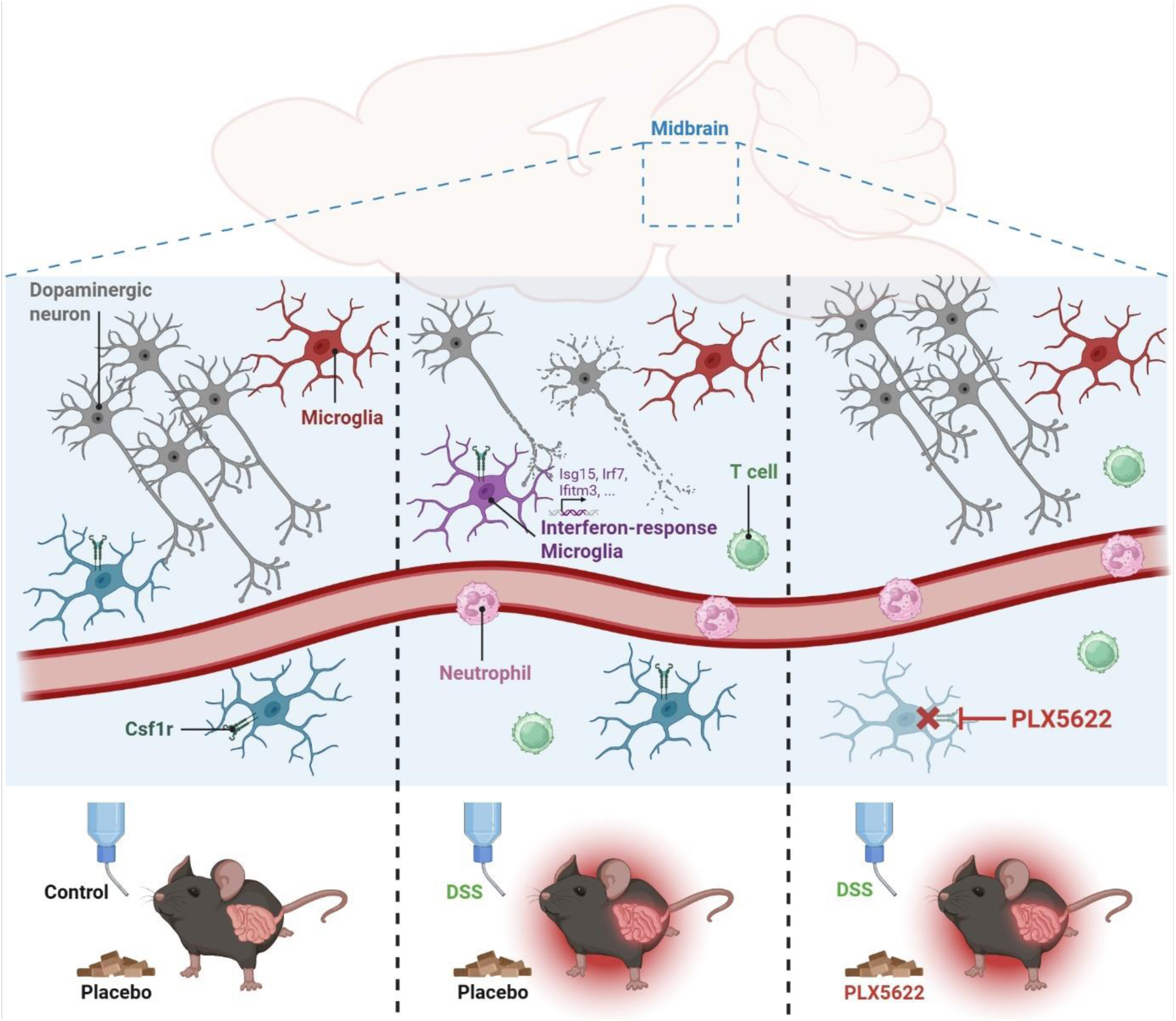

## Introduction

The gut-immune-brain axis describes a vivid immune crosstalk between the gut and the brain and has been proposed to play a fundamental role in numerous brain disorders^[1–3]^. Importantly, inflammatory bowel disease (IBD) is linked to neuroinflammation and neuropsychiatric comorbidity^[4]^, and specifically associated with Parkinson’s Disease (PD) by both genetic and epidemiological evidence^[5]^. Vice versa, PD may be driven by early microbial and inflammatory stimuli originating from the gut^[6, 7]^. However, underlying cellular mechanisms linking chronic intestinal inflammation to PD-related neurodegeneration are not fully understood. A stepwise cascade was proposed, initialized by propagation of inflammatory cues across a leaky gut vascular barrier leading to systemic inflammation^[4]^. This, in turn, can drive neuroinflammation, characterized by the infiltration of blood-derived immune cells into the brain parenchyma and the activation of brain-resident microglia, which are highly responsive to gut-derived stimuli^[8, 9]^. Yet, whether and how gut-induced neuroinflammation can drive neurodegeneration remains to be elucidated. While rodent models of IBD, particularly based on dextran sulfate sodium (DSS) colitis, have demonstrated neuroinflammation^[10–12]^ and suggested a loss of dopaminergic neurons in the midbrain^[11]^, a cellular link between both findings remains elusive. Importantly, the precise cellular composition of the midbrain immune response to chronic colitis has not been resolved yet.

In this study, we hypothesize that adult-onset relapsing-remitting experimental colitis drives PD-related neurodegeneration dependent on microglia. We show that chronic DSS colitis results in a systemic inflammation propagating into the brain, where microglial activation is present in the midbrain, but not the striatum, hippocampus, and cortex. This corresponds to an inflammatory signature in bulk tissue RNA-sequencing and proteomics of the midbrain compared to the striatum. The mapping of the midbrain immune cell response to chronic DSS colitis at the single-cell level reveals an expansion of inflammatory microglia clusters including interferon response microglia, parenchymal CD8^+^ T cell infiltration, and vessel-associated neutrophils. Within this complex midbrain immune response, integrated multi-omic analysis predominantly links microglial clusters to alterations along the gut-immune-midbrain axis during colitis. Finally, we show that myeloid cell targeting using the colony-stimulating factor 1 receptor (Csf1r) antagonist PLX5622 after the first cycle of chronic DSS colitis leads to a significantly reduced number of microglia and prevents the observed inflammation-driven loss of dopaminergic neurons in the substantia nigra pars compacta. Of note, Csf1r inhibition does not ameliorate mucosal pathology or prevent the migration of T cells and neutrophils to the midbrain, indicating a microglia-dependent neuroprotective effect.

Collectively, we provide an integrated multi-omic characterization and single-cell resolution of the neuroinflammatory midbrain response to chronic colitis. Within a complex neuroinflammatory pattern, we identify Csf1r-dependent microglia as a treatment target to protect dopaminergic neurons in chronic gut-derived inflammation, with potential relevance for both IBD and PD.

## Methods

### Animals

Experiments were performed with C57BL/6J mice (Charles River Laboratories, Sulzfeld, Germany; The Jackson Laboratory, Bar Harbor, ME, USA) and *Hexb^tdTomato^* mice^[13]^. Mice were housed in individually ventilated cages under conventional animal housing conditions (12-hour light/dark cycle, 22°C) and were provided sterile water and standard laboratory chow *ad libitum*. All animal experiments were approved by the Government of Lower Franconia, Würzburg, Germany (RUF-55.2.2-2532-2-1638) and were performed in accordance to the EU Directive on the Protection of Animals Used for Scientific Purposes (2010/63/EU) in the preclinical experimental animal center in the Franz-Penzoldt-Center in Erlangen, Germany.

### Colitis induction and scoring

Starting at an age of 20 weeks, chronic colitis was induced in three cycles, each consisting of *ad libitum* provision of 2% (w/v) dextran sodium sulfate (DSS; colitis grade, 36 – 50 kDa, MP Biomedicals) in drinking water for 5 days followed by normal drinking water for 10 days. Control mice received normal drinking water throughout the study period. Disease severity was monitored on a daily basis by scoring changes in bodyweight, grooming and nesting behavior as well as clinical symptoms, which were combined to yield the disease associated index. On day 36 of 45, colonoscopy was performed under continuous anaesthesia with 2% isofluorane and by using a 1.9 mm telescope (Karl Storz, Tuttlingen, Germany) connected to an air pump. Colitis scoring was performed according to the modified murine endoscopic index of colitis severity described by Becker, Fantini ^[14]^.

Additionally, colitis severity was scored by histopathological analysis of hematoxylin and eosin (H&E) stained colon sections as previously described^[15]^. The sum scores for (a) intestinal integrity (0-3), (b) extent of mucosal inflammation (0-3), and (c) submucosal pathological changes (0-3) of each mouse were taken as an indicator for disease severity.

### PLX5622 treatment

Microglia were depleted by feeding 1200 ppm PLX5622 supplemented chow (Hycultec, Beutelsbach, Germany; ssniff Spezialdiäten, Soest, Germany) while control mice were fed ingredient-matched PLX5622-free placebo chow starting at day 11 for 35 days. Mice were randomly distributed into four groups: (1) control mice receiving placebo chow, (2) control mice receiving PLX5622-supplemented chow, (3) chronic DSS colitis mice receiving placebo chow, and (4) chronic DSS colitis mice receiving PLX5622-supplemented chow.

### Tissue collection

Mice were anesthetized with 2% isofluorane and blood was collected by cardiac puncture, followed by immediate transcardiac perfusion using ice-cold phosphate buffered saline (PBS). The colon and intestine were flushed with PBS to remove content and samples from the distal colon and ileum, liver, and lung were either snap-frozen or fixed in 4.5% formaldehyde and embedded in paraffin. The brain was isolated and hemispheres or regions were dissected.

For bulk and single-cell RNA-sequencing experiments, the choroid plexus from the lateral and fourth ventricles was removed before dissecting the midbrain and striatum to prevent sample contamination as previously reported^[16]^. Isolated midbrain and striatum tissue from contralateral brain hemispheres were used for proteome and transcriptome analyses.

### Serum multiplex ELISA

Serum cytokine levels were measured using the V-PLEX Mouse Cytokine 19-Plex Kit (Meso Scale Diagnostics, Rockville, MD, USA) according to the manufacturer’s protocol. Briefly, serum was prepared by centrifugation (2,000 g, 10 min, 4°C) of blood collected by cardiac puncture, and was snap-frozen in liquid nitrogen and stored at -80°C until further analysis. Samples were measured in duplicates at a MESO QuickPlex SQ 120MM (Meso Scale Diagnostics) and analyzed with the software MSD DISCOVERY WORKBENCH® (v4.0, Meso Scale Diagnostics).

### Flow cytometry

Immune cells were isolated from PBS-perfused whole brain tissue as described by Masuda, Amann ^[13]^ with minor modifications. In brief, one brain hemisphere was manually homogenized in ice-cold Hanks’ balanced salt solution containing 15 mM HEPES buffer and 0.54% glucose using a dounce tissue grinder and filtered through a 70 µm cell strainer before centrifugation (200 g, 5 min, 4°C). Cells were separated in 37% Percoll by gradient centrifugation (800 g, 35 min, 4°C, no break), washed in ice-cold PBS and transferred to a 94-well plate. Single-cell suspensions were stained with LIVE/DEAD™ Aqua Dye (Thermo Fisher Scientific, Waltham, MA, USA; 1:1,000 in PBS) for 10 min at 4°C, washed and stained with fluorophore-conjugated surface antibodies diluted in PBS for 30 min at 4°C. Data were acquired on a Cytek Northern Lights (Cytek Biosciences, Fremont, CA, USA) using the SpectroFlo software (v3.3.0, Cytek Biosciences) and analyzed using OMIQ (Dotmatics, Boston, MA, USA). Detailed information on anti-mouse antibodies used for flow cytometry is provided in the Reporting Summary.

### Immunofluorescence stainings

Free-floating immunofluorescence staining was performed as previously described^[15]^. Brain hemispheres and colon tissue were fixed in 4% paraformaldehyde and rehydrated in 30% sucrose solution, followed by serially cutting into 20 µm (brain) sagittal or coronal sections or 40 µm (colon) sections on a Leica SM2010 R sliding microtome (Leica Biosystems, Nussloch, Germany). Antigen retrieval by pre-treatment in citrate buffer (0.1 M citric acid, 0.1 M Tris-sodium citrate in ddH_2_O) for 30 min at 80°C was performed prior to all antibody exposures except CD68. Tissue sections were incubated in blocking solution (3% donkey serum, 0.3% Triton™ X-100 in tris-based saline (TBS)) and stained overnight at 4°C with specific primary antibodies (detailed information provided in the Reporting Summary). Samples were washed with TBS and counterstained with respective fluorophore-conjugated secondary antibodies (Thermo Fisher Scientific). After washing with TBS, cell nuclei were stained using 4’,6-Diamino-2-phenylindole (DAPI) and tissue slices were mounted with ProLong Gold Antifade Mountant (Thermo Fisher Scientific).

### Image acquisition and analysis

Immunofluorescence-stained sections were imaged using either a laser scanning confocal microscope LSM 780 (Carl Zeiss Microscopy, Jena, Germany) on an inverted Axio Observer with a Plan-Apochromat 20x/0.8 M27, a Plan-Apochromat 40/1.46 Oil DIC M27, or alpha Plan-Apochromat 63x/1.46 Oil Korr M27 objective (Carl Zeiss Microscopy). Maximum intensity projections were generated using the Zeiss Zen 2012 (black edition) software (v8.1.0.484), and cells were counted using the Zeiss Zen 2012 (blue edition) software (v1.1.2.0). All analyses were performed blinded to group allocation.

### Analysis of microglia activation

Microglia activation was assessed based on the morphology and expression of the lysosomal protein CD68 as previously described^[15, 17]^. From each animal, four images were acquired per brain region from 2-3 brain sections under a 40x oil objective (scan mode: plane, stack, z: 20 µm, scaling: 0.5 µm, pixel dwell time: 1.54 µs). Scoring was performed on maximum intensity projections covering a total number of Iba1^+^ cells (cDSS/control) in the cortex (277/300), hippocampus (223/277), striatum (294/315), and substantia nigra (436/459). Microglia morphology was investigated both by Sholl analysis and analysis of microglial processes from 3D-reconstructed images in Imaris (v7.6.4, Oxford Instruments, Belfast, UK).

### Quantification of immune cells

Microglia (Iba1^+^), neutrophils (Ly6G^+^), and T cells (CD3^+^) were quantified from 3-4 images per brain region in 2-3 brain sections per mouse under a 40x oil objective (scan mode: plane, stack, z: 20 µm, scaling: 1.0 µm, pixel dwell time: 1.54 µs). Counterstaining with collagen IV allowed to distinguish CD3^+^CD8^+^ T cells inside vessels (vascular), in the perivascular space (perivascular), and in the brain parenchyma (parenchymal). Ly6G^+^ cells adjacent to collagen IV^+^ basal laminae were considered as vessel-associated neutrophils.

Myeloid cells (Iba1^+^) in the colon were quantified in the mucosa, submucosa, and muscularis from 3 tile scan images (2×2) in three colon sections per mouse taken with a 20x objective (scan mode: tile, stack, z: 40 µm, scaling: 2.0 µm, pixel dwell time: 4.10 µs). Counting was performed on maximum intensity projections.

### Quantification of dopaminergic neurons

The total number of dopaminergic neurons in the substantia nigra pars compacta was analyzed from tyrosine hydroxylase (TH) staining on every 12^th^ coronal brain section from each animal. Images were acquired from each brain section covering the total substantia nigra under a 20x objective (scan mode: tile, stack, z: 20 µm, scaling: 1 µm, pixel dwell time: 4.01 µs) and the substantia nigra pars compacta was identified from maximum intensity projections using the Allen Brain Atlas of the mouse brain as reference^[18]^. All TH^+^ cells were manually counted, and the number of cells was multiplied by 12 to determine the total number of TH^+^ cells in the substantia nigra pars compacta of one brain hemisphere.

### Protein extraction and preparation for mass spectrometry

Isolated midbrain and striatum tissue from contralateral brain hemispheres were used for proteome and transcriptome analysis. Total protein was isolated from snap frozen tissue by homogenization in 1:6 w/v homogenization buffer (50 mM Tris/HCl, 150 mM NaCl, pH 8.0) supplemented with 1x cOmplete™ EDTA-free Protease Inhibitor Cocktail (Roche Diagnostics, Mannheim, Germany) and mechanical dissociation using a Potter S Homogenizer (800 U/min; Sartorius Stedim Biotech, Göttingen, Germany). Homogenous samples were centrifuged (10,000 g, 10 min, 4°C) and protein concentration was quantified using the Pierce BCA Protein Assay Kit according to the manufacturers protocol (Thermo Fisher Scientific). Colorimetric absorbance was measured at a CLARIOstar PLUS high-performance microplate reader (BMG Labtech, Ortenberg, Germany). Samples were stored at -80°C until mass spectrometric analysis.

### Mass spectrometry

Protein analysis was performed using the Single-Pot Solid-Phase-enhanced Sample Preparation method^[19, 20]^. In brief, 20 µg protein were mixed with Sera-Mag™ SpeedBeads A and B (Cytiva, formerly GE Healthcare Life Sciences, Marlborough, MA, USA) and binding was initiated by adding absolute ethanol. After incubation, beads were washed with 80 % ethanol and resuspended in 100 mM ammonium acetate buffer at a Hamilton Microlab Prep automated liquid handling platform (Hamilton Company, Reno, NV, USA) before on-bead digestion with trypsin. Resulting peptides were eluted in 1 % formic acid and stored at -20°C until measurement. Samples were measured on an Orbitrap Eclipse Tribrid Mass Spectrometer coupled to an UltiMate 3000 Nano-High-performance liquid chromatography (Thermo Fisher Scientific) via a Nanospray Flex Ion Source equipped with a Sonation column oven and front-end High Field Asymmetric Waveform Ion Mobility Spectrometry interface. Raw files were processed using the ProteoWizard software^[21]^ and proteins were identified using the data independent acquisition neural network software^[22]^. Data was further processed using the software Perseus (v2.0.11)^[23]^ and visualized using the ggplot2 package built in R^[24]^.

### RNA isolation and reverse transcription-quantitative PCR

RNA for reverse transcription-quantitative PCR (RT-qPCR) was isolated from fresh-frozen tissue using the RNeasy Mini Kit (Qiagen, Hilden, Germany) and reversely transcribed using the GoScript™ Reverse Transcription System (Qiagen) according to the manufacturer’s protocols. Gene expression on mRNA level was assessed using the SsoFast™ EvaGreen® Supermix (Bio-Rad, Hercules, CA, USA) according to the manufacturer’s protocol and samples were measured at a LightCycler® 96 System (Roche Diagnostics) and analyzed using the LightCycler® 96 Software (v1.10.1320, Roche Diagnostics). Primer sequences used for RT-qPCR are listed in Table S1.

### Bulk tissue RNA-sequencing

Midbrain and striatum tissue were dissected from PBS-perfused brains on ice and homogenized in QIAzol™ Lysis Reagent. Libraries were prepared as previously described^[8]^. In brief, total RNA was isolated using the Direct-zol RNA Microprep Kit (Zymo Research Europe, Heilbronn, Germany) and polyadenylated mRNA was enriched by incubation with Oligo d(T)_25_ Magnetic Beads (New England Biolabs, Ipswich, MA, USA). The SuperScript™ III First-Strand Synthesis SuperMix (Thermo Fisher Scientific) was used for first-strand cDNA synthesis and PCR products were purified using RNAClean XP beads according to the manufacturer’s instruction (Beckman Coulter Life Sciences, Indianapolis, IN, USA). The RNA/cDNA double-stranded hybrid was labeled with dNTP/dUTP and dsDNA was purified using Sera-Mag™ SpeedBeads B (GE Healthcare Life Sciences). The purified dsDNA underwent end repair by blunting, A-tailing, and adaptor ligation using NEXTflex™ DNA Barcodes (Revvity, Waltham, MA, USA) as previously described^[25]^. The libraries were PCR amplified for 16 cycles, selected for fragments (size 200-500 bp) by gel extraction, and were quantified using a Qubit dsDNA High Sensitivity Assay Kit (Thermo Fisher Scientific), and sequenced on a NovaSeq 6000 for 51 cycles according to the manufacturer’s instructions (Illumina, San Diego, CA, USA).

RNA sequencing reads from FASTQ files were mapped to the mus musculus reference genome (mm10) using Spliced Transcript Alignment to a Reference (STAR, v2.7.9a) with default parameters and further analyzed using HOMER (v5.1) as previously described^[26, 27]^. Differential gene expression was assessed using DESeq2 (v1.48.2) and defined by adjusted p-value < 0.05 and log_2_ fold change > 0.5 or < -0.5^[28]^. Data normalization and visualization was done using R. Pathway analyses were performed using clusterProfiler (v4.16.0)^[29]^.

### Single-cell RNA-sequencing

Immune cells were isolated from midbrain tissue by enzymatic digestion as described by Scheyltjens, Van Hove ^[30]^ with minor modifications. The transcriptional inhibitor actinomycin D (ActD) was added in decreasing concentrations to media and buffers to prevent induction of immediate early genes during the dissociation process^[31]^. Briefly, midbrain tissue isolated from PBS-perfused brains was cut in 1-2 mm pieces in collection buffer (30 µM ActD in RPMI) and incubated with enzyme stock solution (90 U/ml DNase, 30 U/ml Collagenase I, 1200 U/ml Collagenase IV, 15 µM ActD in 1x HBSS) for 35 min (37°C, 300 rpm). Cell homogenates were filtered through a 100 µm mesh filter, pooling tissue from two mice. Cells were separated by gradient centrifugation using 37 % Percoll (800 g, 35 min, 4°C, no acceleration, no deceleration) and washed in FACS buffer (5 mM EDTA, 2 % BSA in 1x HBSS) before staining for CD11b and CD45 (20 min, 4°C). Cells were washed in FACS buffer and pooled, resulting in one sample per group. CD45-positive cells were sorted into 1x PBS on a MoFlo™ XDP flow cytometer (Beckman Coulter) in the Core Unit Cell Sorting and Immunomonitoring of the Medical Faculty, Friedrich-Alexander-Universität Erlangen-Nürnberg, Erlangen, Germany. The yield of CD45-positive cells sorted from pooled tissue of 6 mice, was 221,880 (Control) and 180,660 (cDSS). The cell concentration of each sample was adjusted to 1000 cells/µl using 1x PBS. Libraries were prepared using the Chromium Next GEM Single Cell 3’ Reagent Kits v3.1 CG000315 Rev E according to the manufacturer’s protocol (10x Genomics) with a targeted cell recovery of 10,000 cells. The Chromium Single Cell 3’ Gene Expression Dual Index library was sequenced using a NovaSeq 6000 instrument (Illumina) at Genewiz, Leipzig, Germany.

RNA sequencing reads from FASTQ files were mapped to the mus musculus reference genome (mm10) using the 10x Genomics Software Cell Ranger (10x Genomics, Pleasanton, CA, USA). Subsequent analysis was performed in R (v4.5.1). Quality control, data normalization and scaling, and subsequent clustering was performed using Seurat (v5.3.0)^[32]^. The median was calculated for nCount_RNA, nFeature_RNA and percent.mt and 1.5 times the interquartile range from the first and third quartile were taken as thresholds. In detail, cells with > 500 and < 7500 nFeature_RNA, > 1000 and < 30000 nCount_RNA and < 5% of the detected molecular identifiers mapped to mitochondrial genes were included, resulting in 7070 cells (cDSS) and 6591 cells (Control) post filtering. Principal component analysis (PCA) was performed on the top 2000 highly variable genes prior to clustering using clusterProfiler^[29]^. Clusters were visualized using uniform manifold approximation and projection for dimension reduction (UMAP) and annotated based on the top 10 identified marker genes per cluster using the FindAllMarkers function in Seurat. Pathway analyses were performed using clusterProfiler. Microglia subclusters were named according to previously published guidelines for microglia cell state annotation^[33]^.

### Multi-Omics Factor Analysis (MOFA)

An integrated analysis of different data modalities (bulk RNA-sequencing and proteomics of midbrain tissue, qPCR data of genes expressed in the colon, ELISA of serum cytokines) was conducted using the MOFA2 R package (v1.18.0)^[34]^. Cytokine concentrations were log2 transformed and qPCR data were scaled prior to model training. The identified latent factors were correlated with the 50 top markers per cell type from midbrain single-cell RNA-sequencing data (cell-type signatures) to identify cell types enriched per factor. Fast Gene Set Enrichment Analysis (FGSEA) was conducted to identify cell-type signatures associated with MOFA factors using the FGSEA R package (v1.34.2)^[35]^. Gene weights were extracted from the ‘Transcriptomics midbrain’ view of the trained MOFA+ model and ranked according to their factor loadings. Cell type specific gene signatures were tested for enrichment against this pre-ranked gene list. Normalized enrichment scores and Benjamini-Hochberg adjusted p-values were used to assess significance and correct for multiple testing.

## Statistical analysis

Statistical analyses were performed using R (v4.5.1), RStudio (v2025.09.1), or Prism 10 (v10.2.0). All data are shown as mean ± standard deviation (s.d.). Statistical test performed for individual data analysis are indicated in the respective figure legends.

## Data availability

Bulk proteomics data have been deposited at the Proteomics IDEntifications Database (PRIDE) and are accessible under the accession number PXD069675 upon publication. Bulk and single-cell RNA-sequencing data for this study have been deposited in the European Nucleotide Archive (ENA) at EMBL-EBI under the accession number PRJEB100827 (https://www.ebi.ac.uk/ena/browser/view/PRJEB100827) and will be made accessible upon publication.

The authors declare that all other data supporting the findings of this study as well as source data are provided with this paper and its supplementary information files.

## Code availability

All codes used in this study can be provided upon reasonable request.

## Results

### Cyclic DSS treatment induces chronic relapsing-remitting colitis accompanied by systemic inflammation

To induce chronic relapsing-remitting colitis, we treated male and female C57BL/6J mice aged 20 weeks with DSS in three cycles, each consisting of 5 days of DSS + 10 days of recovery (Fig. 1A, Fig. S1A). Colitis induction by DSS ingestion was confirmed by increased mucosal granularity, adhesive stool residues, thickened, non-translucent colon walls, and fibrin deposition observed by colonoscopy on day 36, quantified by the increased colonoscopy score in chronic DSS colitis mice (Fig. S1B). Post dissection, the DSS group showed a significant shortening of colon length (Fig. S1C). An increased expression of *Il1b*, *Cxcl13*, and *Tnf* in the colon, but not in the small intestine was measured by reverse transcription-quantitative PCR (RT-qPCR), confirming the spatial nature of DSS-mediated induction of inflammation within the large intestine (Fig. S1D).

**Figure 1.**
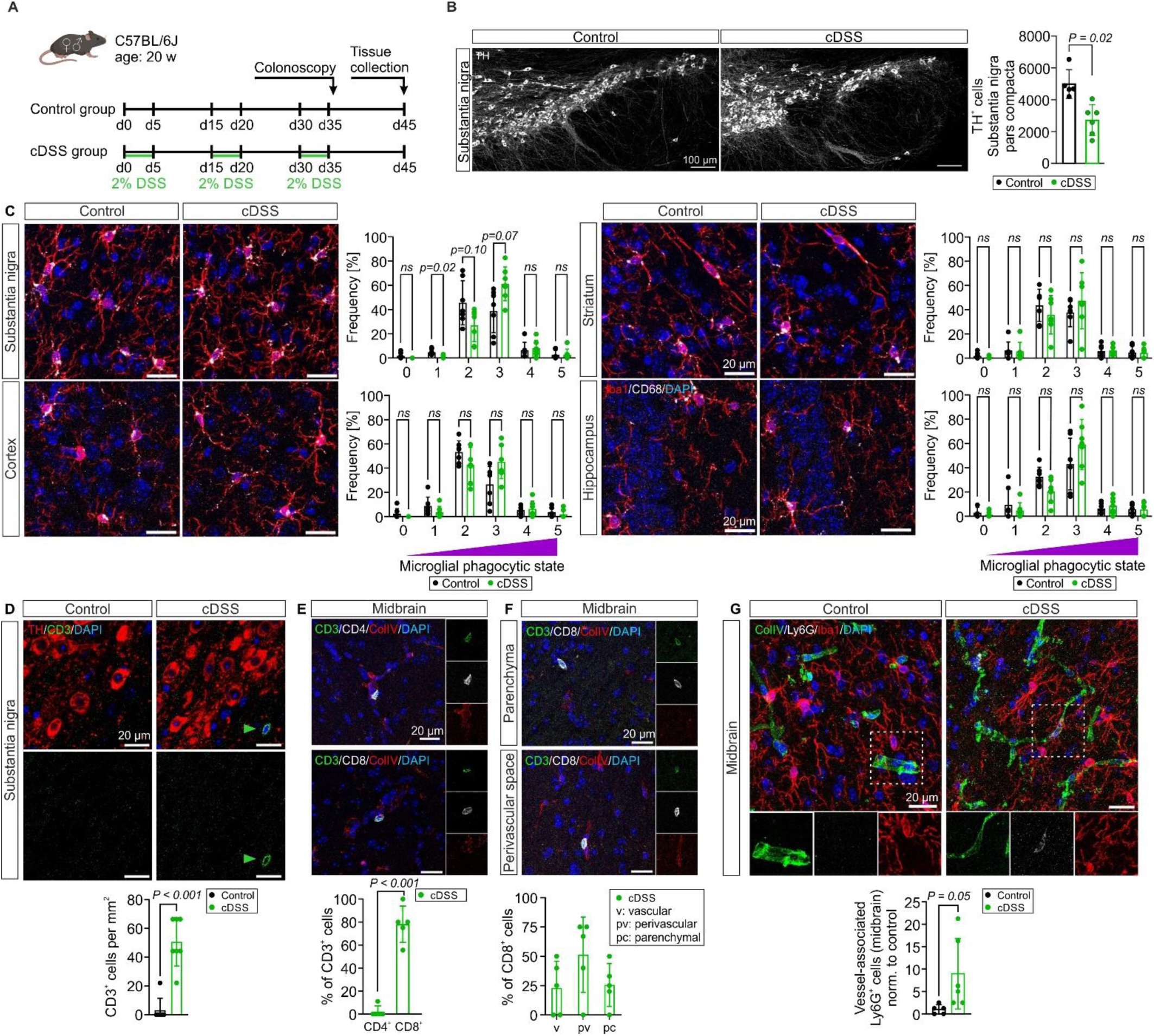
Chronic experimental colitis causes dopaminergic neuroinflammation and a cellular immune response in the midbrain. **A** Timeline of colitis induction in male and female C57BL/6 mice aged 20 weeks at the start of the experiment. **B** Immunostaining for tyrosine hydroxylase (TH, white) and quantification of total number of dopaminergic neurons in the substantia nigra pars compacta (*n* = 5-6 mice per group, unpaired *t*-test). Scale bars, 100 µm. **C** Immunostaining for Iba1 (red) and CD68 (white) in C57BL/6J mice and proportion of microglia phagocytic states in the substantia nigra (top left), the striatum (top right), the cortex (bottom left), and the hippocampus (bottom right) (*n* = 7 mice per group, total 22-77 microglia in each region; multiple unpaired *t*-tests). Scale bars, 20 µm. **D** Immunostaining for CD3 (green) and TH (red) in C57BL/6J mice (top). Arrowheads indicate CD3^+^ T cells in the substantia nigra. Quantification of CD3^+^ T cell density in the substantia nigra (bottom; *n* = 7 mice per group). Scale bars, 20 µm. **E** Immunostaining for CD3 (green), CD4 (white, top), CD8 (white, bottom) and ColIV (red) in the midbrain (top) in chronic DSS C57BL/6J mice. Proportion of CD4^+^ and CD8^+^ T cells within the total CD3^+^ T cell population in the midbrain (bottom; *n* = 5 mice). Scale bars, 20 µm. **F** Immunostaining for CD3 (green), CD8 (white), and ColIV (red) in the midbrain parenchyma (top) as well as perivascular space (middle) in chronic DSS C57BL/6J mice. Proportion of vascular, perivascular and parenchymal CD8^+^ T cells within all CD8^+^ T cells in the midbrain (bottom; *n* = 5 mice). Scale bars, 20 µm. **G** Immunostaining for Ly6G (white), Iba1 (red), and ColIV (green) in C57BL/6J mice (top). Proportion of vessel-associated neutrophils in the midbrain normalized to control (bottom; *n* = 5-6 mice per group). Scale bars, 20 µm. Two-tailed, unpaired *t*-test, if not otherwise indicated. Data are presented as mean ± s.d. Icons in **A** were created using BioRender.com.

Next, we investigated systemic inflammation in peripheral organs and the blood during chronic DSS colitis as previously reported^[12, 36]^.This is considered as an important prerequisite for inflammatory propagation from the gut into the CNS^[4]^. Indeed, we found signs of extraintestinal inflammation during the chronic phase of experimental colitis, indicated by an increased spleen length and relative weight in proportion to bodyweight (Fig. S1E, F). In addition, we detected a significantly higher expression of *Il1b* and *Tnf* in the lung, and of *Cxcl13* and *Tnf* in the liver by RT-qPCR analyses (Fig. S1G). Moreover, increased serum levels of the inflammation-associated cytokines IFNγ, TNFα, IL-1β, IL-2, IL-6, IL-10, and CXCL10 confirmed a systemic immune cell activation as a response to chronic mucosal inflammation (Fig. S1H). Serum levels of the CXCL2 were reduced in chronic DSS colitis mice, while comparable levels between mice with chronic DSS colitis and controls were measured for IL-4, IL-5, IL-27p28/IL-30, IL-33, CCL2, CCL3, and CXCL1 (Fig. S1H). In sum, cyclic DSS administration resulted in a moderate relapsing-remitting colitis associated with systemic inflammation.

### Chronic colitis causes dopaminergic neuron loss and neuroinflammation in the midbrain

Given the link of IBD to PD, we next investigated whether chronic experimental colitis causes PD-related neurodegeneration. Indeed, we observed a significant loss of tyrosine hydroxylase (TH) expressing dopaminergic neurons in the substantia nigra pars compacta during chronic DSS colitis (Fig. 1B). To unravel the underlying cellular immune mechanism behind this neuronal loss, we next aimed to characterize the CNS immune landscape during with a particular focus on parenchymal microglia. We therefore induced chronic colitis in the microglia-specific *Hexb^tdTomato^* mouse model^[13]^ to distinguish *tdTomato^+^* microglia from *tdTomato^-^* infiltrating monocyte-derived macrophages and to define their responses to chronic colitis (Fig. S2A). *Hexb^tdTomato^* mice showed similar pathophysiological features in response to chronic colitis compared to non-transgenic C57BL/6J mice, including increased colonoscopy scores, reduced colon length, and increased spleen length and weight (Fig. S2B-E). We first investigated the cellular CNS immune response in whole brain tissue of *Hexb^tdTomato^*mice by flow cytometry (Fig. S2F-H, Fig. S3). While we found similar levels of CD45^+^ immune cells in the brain of chronic DSS colitis and control mice (Fig. S2F), we detect a decreased percentage of CD45^int^CD11b^+^ microglia and increased percentages of T cells, B cells, and neutrophils of all CD45^+^ immune cells in whole brain homogenates of chronic DSS colitis mice (Fig. S2G). Notably, the percentages of CD4^+^ T cells, CD8^+^ T cells, and double negative CD4^-^CD8^-^ T cells were all higher in chronic DSS colitis mice (Fig. S2H). As we determined similar densities of Iba1^+^ cells in the cortex, hippocampus, striatum, and substantia nigra by regional immunofluorescence analysis (Fig. S4A), the relative decrease in the microglial frequency determined by flow cytometry (Fig. S2G) rather results from peripheral immune cell infiltration than from a decrease of cell numbers within the microglial compartment. The identity of Iba1^+^ cells as microglia was confirmed based on tdTomato-expression using immunofluorescence and flow cytometry of brain tissue of *Hexb^tdTomato^* mice with chronic colitis (Fig. S2I-K). Semi-quantitative scoring of microglial cells based on morphology and CD68 expression as previously described^[15, 17]^ indicated lower numbers of resting microglia (score 1) and a tendency towards increased numbers of moderately phagocytic microglia (score 3) in the substantia nigra, but not in the striatum, hippocampus, and cortex in chronic DSS colitis mice (Fig. 1C). As microglial morphology did not differ between chronic colitis and controls based on Sholl analysis and other morphometric measures (Fig. S4B, C), these changes were mainly related to CD68 expression. To determine whether the increase in T cells and granulocytes found by flow cytometry was also prominent in the midbrain and specifically the substantia nigra, we performed immunofluorescence staining together with collagen IV to localize peripheral immune cells to the vascular, perivascular or parenchymal compartment. We observed significantly increased numbers of CD3^+^ T cells in the substantia nigra of mice with chronic DSS colitis (Fig. 1D), primarily accounted for by CD8+ cytotoxic T cells (Fig. 1E). Co-staining with the basement membrane marker collagen IV showed parenchymal localization of some CD8^+^ T cells, though most CD8^+^ T cells were localized inside vessels or in the perivascular space (Fig. 1F). The increased numbers of neutrophils previously detected in flow cytometry of the whole brain were exclusively located within close proximity to blood vessels, but not in the midbrain parenchyma of chronic DSS colitis mice (Fig. 1G).

In conclusion, relapsing-remitting colitis induced by cyclic DSS intake led to a mixed innate and adaptive cellular immune response in the brain, which was pronounced in the midbrain. The diverse neuroimmune response in the midbrain included parenchymal infiltration of CD8^+^ T cells, neutrophil adhesion to brain vessels, and propagation of microglia towards an activated phagocytic phenotype.

### The midbrain-specific transcriptional and proteomic inflammatory signature in chronic colitis

Next, we more precisely characterized the regional CNS response to chronic colitis by bulk RNA-sequencing of the midbrain and striatum alongside proteomic analysis of the contralateral side (Fig. 2A). We observed 129 significantly differentially expressed genes (DEGs) in the midbrain in chronic DSS colitis mice compared to littermate controls (Fig. 2B, C). The 79 upregulated genes were associated with complement activation, humoral immune response, and synapse pruning, whereas the 50 downregulated genes were linked to nervous system development, neurogenesis, and synapse assembly (Fig. 2D). KEGG pathway enrichment analysis identified an association with pathways of neurodegeneration (Fig. 2E). We confirmed the significantly higher expression of microglia-related genes *CD68* and *C1qa, C1qb, and C1qc* in the midbrain of chronic DSS colitis mice by immunofluorescence (Fig. 1C, 2F).

**Figure 2.**
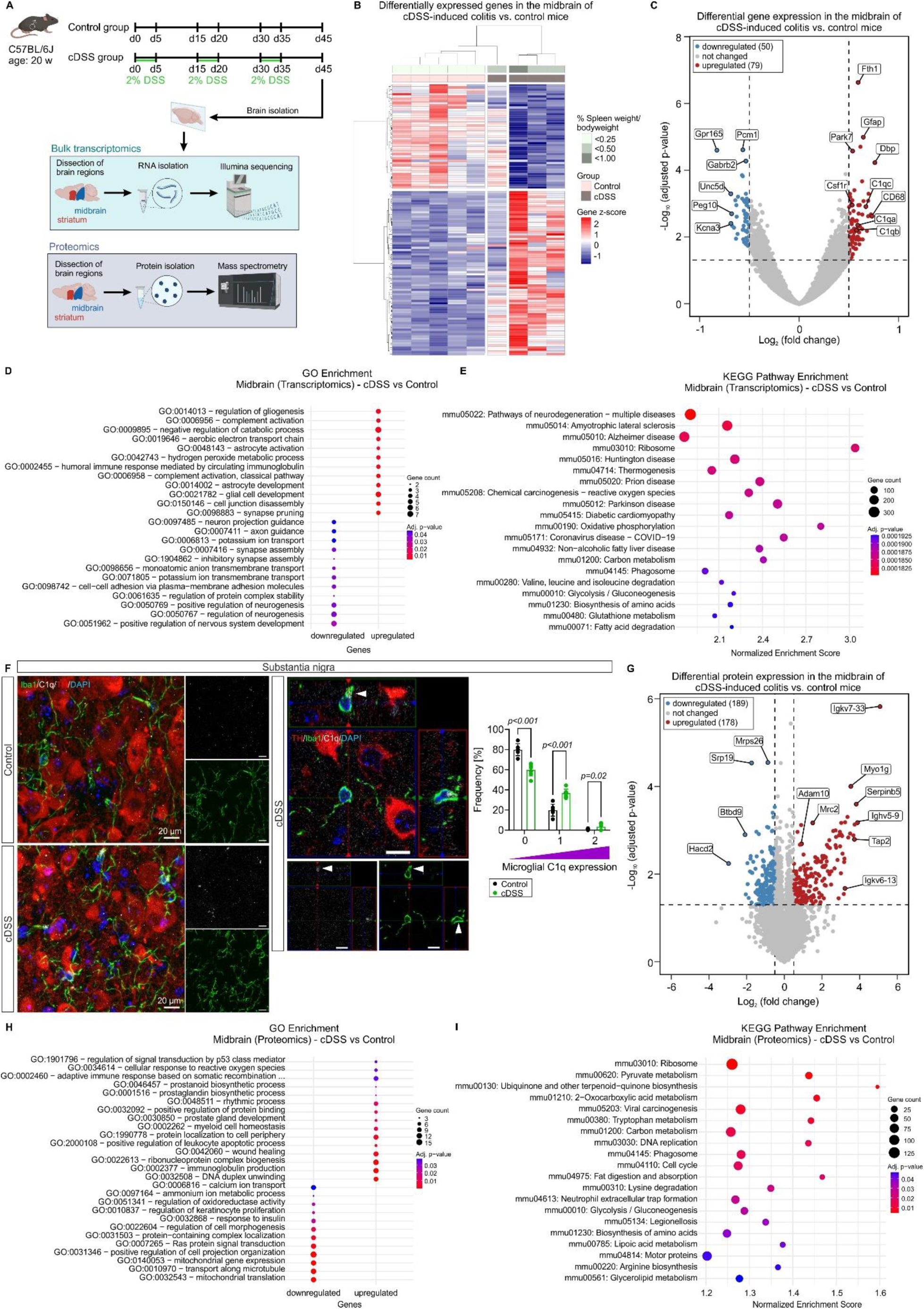
Transcriptome and proteome analyses reveal an inflammatory response in the midbrain. **A** Timeline of colitis induction in male C57BL/6J mice aged 20 weeks at the start of the experiment and scheme representing dissection and proteomic and bulk transcriptomic analysis of midbrain and striatum tissue of contralateral brain hemispheres. **B** Heatmap of differentially expressed genes in the midbrain of chronic DSS colitis vs. control mice. **C** Volcano plot of genes differentially expressed in the midbrain of chronic DSS colitis vs. control mice with the log2 fold change on the x-axis and the -log10 adjusted p-value on the y-axis. **D** Gene Ontology Biological Process analysis of genes significantly higher expressed in the midbrain of chronic DSS colitis mice. **E** KEGG (GSEA) pathway enrichment of genes differentially expressed in the midbrain of chronic DSS colitis mice vs. control mice. **F** Immunostaining for Iba1 (green), C1q (white), TH (red) in brain tissue of C57BL/6J mice (left) and orthogonal view (middle) and quantification of C1q signal (0 = no, 1 = weak, 2 = strong) in Iba1^+^ cells in the substantia nigra (right; *n* = 6-7 mice, multiple unpaired *t*-tests). Scale bars, 20 µm. **G** Volcano plot of proteins differentially expressed in the midbrain of chronic DSS colitis vs. control mice with the log2 fold change on the x-axis and the -log10 adjusted p-value on the y-axis. **H** Gene Ontology Biological Process analysis of proteins significantly higher expressed in the midbrain of chronic DSS colitis mice. **I** KEGG (GSEA) pathway enrichment of proteins differentially expressed in the midbrain of chronic DSS colitis mice vs. control mice. Data in **F** are presented as mean ± s.d, and each point represents the mean value of one mouse. Icons in **A** were created with BioRender.com.

In line with transcriptional alterations, proteomics analysis revealed 367 proteins differentially expressed in the midbrain of chronic DSS colitis mice compared to control mice (Fig. 2G). Pathways associated with the 178 upregulated proteins included cellular response to reactive oxygen species and adaptive immune response, whereas the 189 downregulated proteins were linked to mitochondrial gene expression and cell morphogenesis (Fig. 2H). KEGG pathway enrichment revealed an association with multiple pathways including tryptophan metabolism, phagosome, and neutrophil extracellular trap formation (Fig. 2I). In contrast to the prominent transcriptional and translational changes in the midbrain, gene and protein expression were largely unchanged in the striatum, revealing only two upregulated genes and one downregulated gene as well as 30 upregulated and 42 downregulated proteins in chronic DSS colitis vs. control mice (Fig. S5A-D).

In summary, we found clear evidence for a regional susceptibility of the midbrain to chronic colitis-induced systemic inflammation on gene and protein level.

### Chronic colitis causes a complex cellular immune response including disease-associated microglia in the midbrain

To contextualize the inflammatory to chronic colitis-induced systemic inflammation in the midbrain and to dissect alterations in the cellular composition of midbrain microglia, we sorted CD45^+^ immune cells from the midbrain of mice with chronic DSS colitis and controls and performed single-cell RNA-sequencing (Fig. 3A). After quality control, we conducted uniform manifold approximation and projection (UMAP) analyses from 7070 and 6591 CD45^+^ cells from the midbrain of chronic DSS colitis and control mice, respectively (Fig. 3B, Fig. S6A). Based on gene marker analysis using FindMarkers in Seurat, we identified six microglial clusters (Clusters 0, 1, 2, 3, 4, 10), B cells (Cluster 9), T cells (Cluster 6), NK/NK T cells (Cluster 11), neutrophils (Cluster 5), monocytes (Cluster 8), and CNS-associated macrophages (CAMs)/dendritic cells (Cluster 7, Fig. 3C-F). Compared to the homeostatic midbrain of control mice, non-microglial immune cells were represented at higher levels in the midbrain of chronic DSS mice (cDSS vs control: 24.69 % vs 5.90 %, thereof B cells (2.76 % vs 0.50 %), T cells (4.50 % vs 1.47 %), NK/NK T cells (1.22 % vs 0.56 %), monocytes (2.60 % vs 1.00 %), neutrophils (5.77 % vs 0.56 %), and CAMs/Dendritic cells (1.47 % vs 1.04 %) while the relative proportion of microglia was lower (75.31 % vs 94.10 %), pointing towards a complex multicellular inflammatory response in the midbrain (Fig. 3D). Interestingly, some bone marrow-derived immune cell clusters predominantly found in mice with chronic DSS colitis expressed adhesion and migration molecules such as *Itgal*, *Itga4,* and *Cd44* as well as the Peyer patches-specific homing receptor *Itgb7* (Fig. S6B), indicating their priming in the gut and capacity to subsequently migrate into the CNS. The joint cluster of CAMs and dendritic cells could be segregated by expression of *Mrc1* and *CD209a*, respectively (Fig. S6B). In line with our immunohistological findings, most T cells were CD8^+^ and showed high expression of *Nkg7* and *Ccl5*. To a smaller extent, CD4^+^ cells were also present in the midbrain of chronic DSS colitis mice (Fig. S6B). Sub-analysis of microglia revealed differential representation of six clusters in chronic DSS colitis and control mice, which were classified based on Gene Ontology Biological Process enrichment and previously published microglia cluster defining genes^[33]^ (Fig. 4A-D). The homeostatic microglial clusters 0 and 2 were lower represented in chronic DSS colitis mice (Fig. 4A; cDSS vs control, Cluster 0: 33.46 % vs 37.82 %, Cluster 2: 15.52 % vs 19.31 %). They were characterized by high expression of the established homeostatic microglial markers *Hexb* and *Tmem119* and linked to oxidative phosphorylation (Fig. 4C, D). Similarly, Cluster 1 was associated with homeostatic marker expression and the Gene Ontology terms developmental cell growth and transforming growth factor beta receptor superfamily signaling pathway, but was more strongly represented in control mice (Fig. 4A, C, D; cDSS vs control, Cluster 1: 38.10 % vs 31.28 %). Clusters 3 and 4 were equally represented in mice with chronic colitis and controls, and linked to inflammation related terms (Fig. 4A, D; cDSS vs control, Cluster 3: 6.31 % vs 6.27 %, Cluster 4: 4.11 % vs 3.95 %). Cluster 3 was characterized by increased expression of *Mertk* and reduced expression of *Csf1r* compared to the other microglial clusters (Fig. 4C). *Ccl3* and *CCl4* were specifically enriched in Cluster 4 (Fig. 4A-C). Interestingly, Cluster 10 was strongly induced in mice with chronic DSS colitis and identified as an interferon response microglia cluster with specific expression of *Ifit3*, *Isg15*, and *Irf7* (Fig. 4A-D; cDSS vs control, Cluster 10: 2.50 % vs 1.37 %).

**Figure 3.**
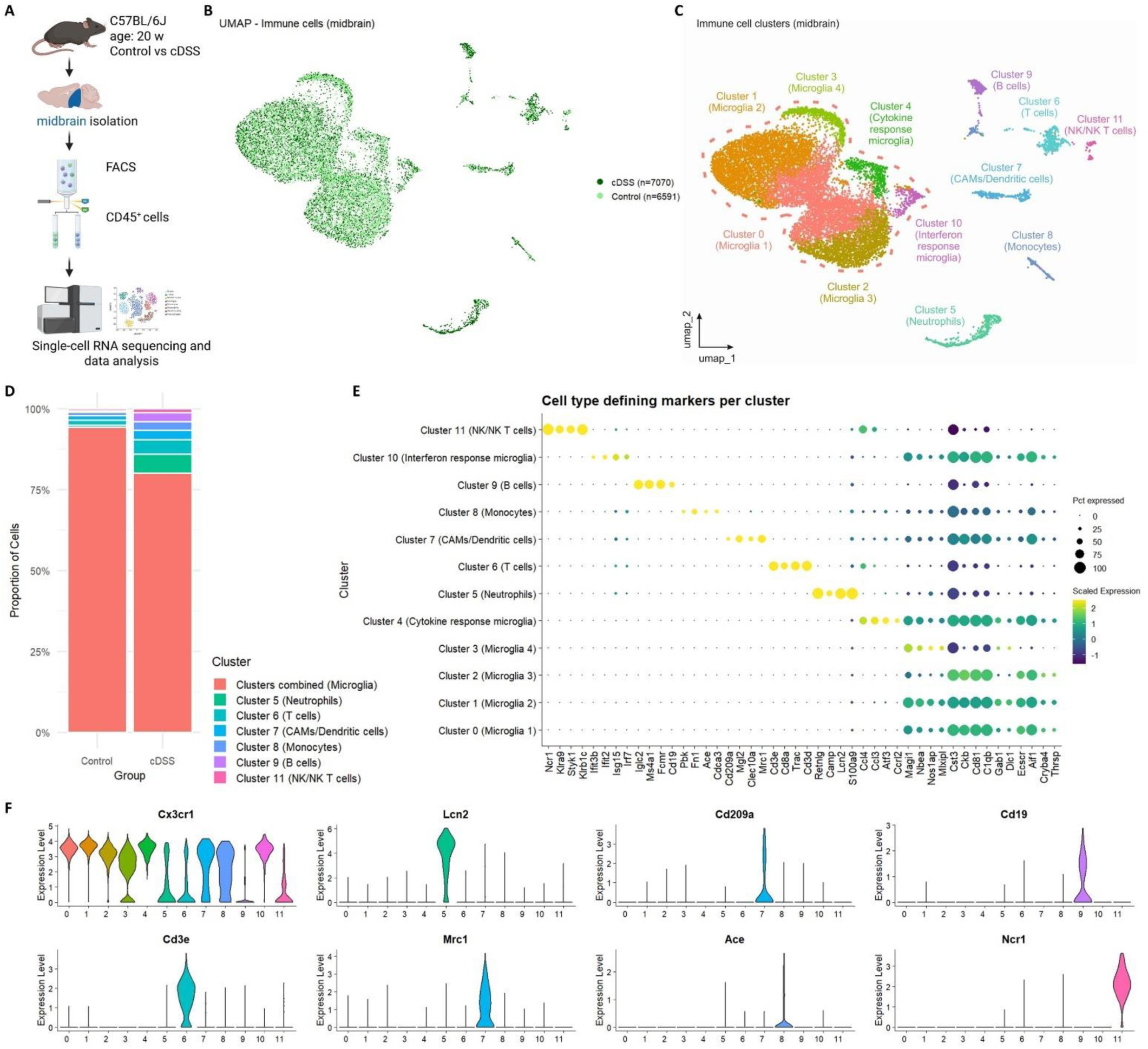
Single-cell RNA-sequencing identifies a complex immune cell response in the midbrain of chronic DSS colitis mice. **A** Experimental design for single-cell RNA-sequencing on sorted CD45^+^ cells from isolated midbrain tissue of male C57BL/6J mice aged 20 weeks at the start of the experiment (n=7 mice pooled per group). **B** UMAP of immune cells split by group (6591 cells control, 7070 cells cDSS). **C** Immune cell clusters identified using the clusterProfiler package in R. **D** Proportion of immune cell clusters of all sorted cells. **E** Dotplot of cell type defining markers per cluster with dots representing the percentage of cells expressing the respective marker (size) and scaled expression (color) in each cluster. **F** Violin plots visualizing expression of the general myeloid cell marker Cx3cr1 and cell type defining markers (Lcn2, Cd209a, Cd19, Cd3e, Mrc1, Ace, Ncr1) in each cluster. Icons in **A** were created with BioRender.com.

**Figure 4.**
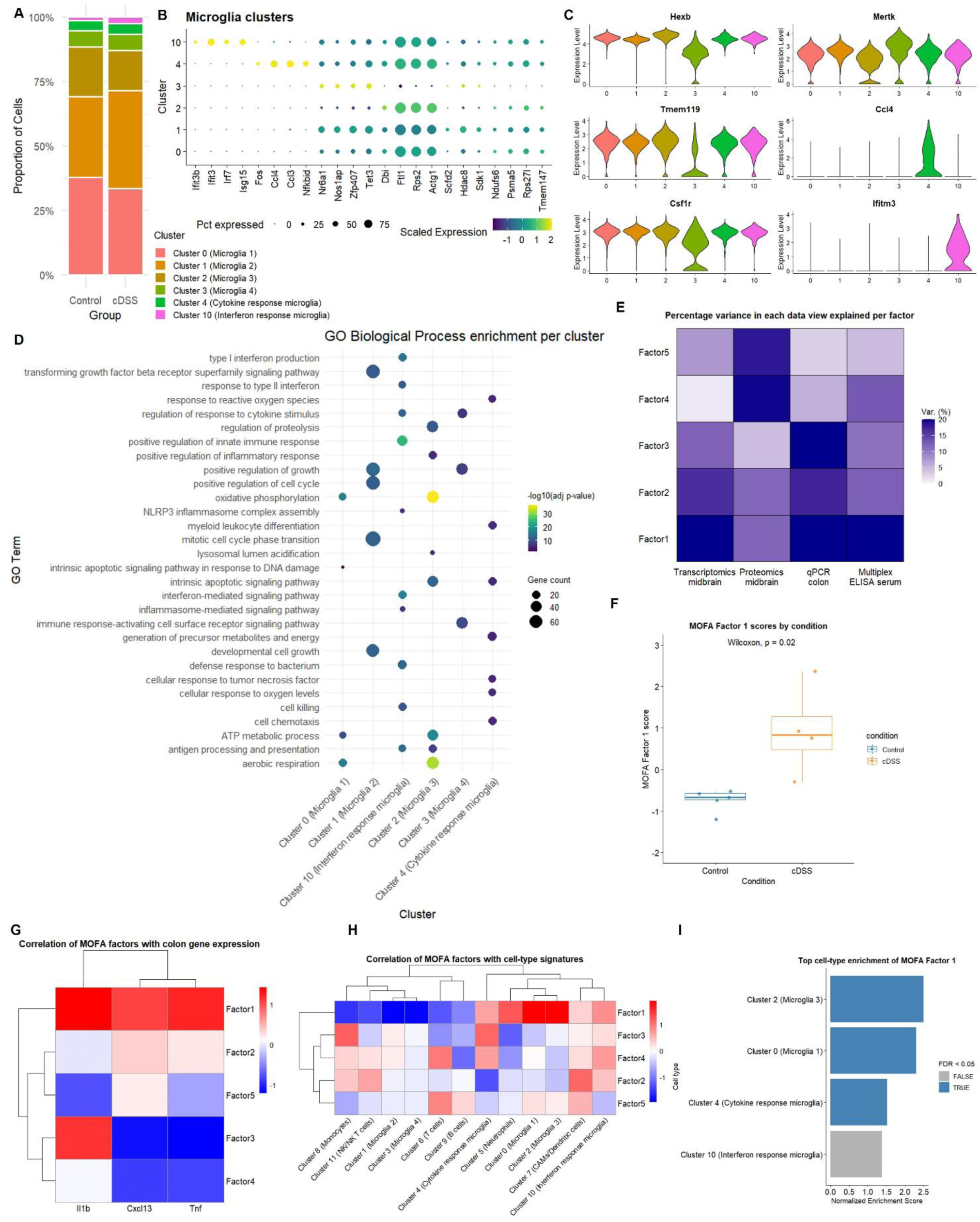
Microglial clusters undergo an inflammatory shift and drive molecular midbrain responses during chronic experimental colitis. **A** Relative abundance of the microglia clusters. **B** Dotplot of microglia cluster defining markers per microglia cluster with dots representing the percentage of cells expressing the respective marker (size) and scaled expression (color) in each cluster. **C** Violin plots visualizing microglial homeostasis marker (Hexb, Tmem119, Csf1r) and activation marker (Mertk, Ccl4, Ifitm3). **D** Gene Ontology Biological Process analysis of genes enriched in each microglia cluster in comparison to all other microglia clusters. **E** MOFA+. Heatmap showing the percentage of variance in each data view explained per factor of the MOFA+ model with 5 factors. **F** Boxplot of MOFA+ Factor 1 scores by condition (*n* = 4-5 mice per group, Wilcoxon signed rank test). **G** Heatmap visualizing the correlation of MOFA factors with the normalized expression of inflammation-associated genes in the colon. **H** Heatmap showing the correlation of each MOFA+ factor with cell type specific gene expression profiles defined from midbrain single-cell RNA sequencing data (cell-type signatures). **I** Bar graph depicting the top four cell types enriched for MOFA+ Factor 1, ranked by the normalized enrichment score. Statistically significant enrichments (FDR < 0.05) are highlighted in blue.

Taken together, chronic colitis-induced systemic inflammation leads to complex alterations in the midbrain immune cell composition characterized by parenchymal infiltration of CD8^+^ T cells, vascular adhesion of neutrophils, and a shift of microglia subpopulations from homeostatic to disease-associated subtypes, including interferon response microglia.

### Multi-Omics Factor Analysis links microglia to the midbrain fingerprint in colitis

Having dissected a complex midbrain immune response to chronic colitis, we next aimed to identify major cellular drivers of midbrain signatures in colitis as potential therapeutic targets to alleviate dopaminergic neuron loss. To identify major sources of cross-modal variation in our multi-omics dataset along the gut-immune-brain axis and to relate these factors to disease state and midbrain immune cells, we applied a comprehensive preprocessing, integration, and harmonization workflow across all data modalities and performed Multi-Omics Factor Analysis v2 (MOFA+). Data of multiple types (views) were integrated, including (1) midbrain bulk RNA sequencing (RNA) and (2) midbrain proteomics (Proteomics) data, (3) qPCR analysis of inflammation-associated genes expressed in the colon (qPCR_colon), and (4) circulating cytokines (ELISA) (Fig. 4 E). MOFA+ captured both shared and modality-specific variation across the multi-omics dataset. Importantly, MOFA+ Factor 1 explained shared variance across all data modalities, indicating a disease-associated pattern. Factor 1 scores were significantly higher in the chronic DSS colitis group compared to controls (Fig. 4F, Wilcoxon rank-sum test, *p = 0.02*) and positively correlated with the expression of inflammation-associated genes, i.e. *Il1b*, *Cxcl13*, *Tnf* (Spearman correlation: *Il1b* r = 0.717, *Cxcl13* r = 0.533, *Tnf* r = 0.617) in the colon (Fig. 4F, G). To assess the cellular basis of this latent program, we correlated MOFA+ factors with cell-type specific expression signatures derived from midbrain single-cell RNA sequencing data (cell-type signatures, Fig. 4H). Of note, MOFA+ Factor 1 scores positively correlated with multiple microglia clusters (Spearman correlation: Cluster 0 r = 0.950, Cluster 2 r = 0.983, Cluster 4 r = 0.683, Cluster 10 r = 0.450), suggesting that this latent factor captures microglia-related variation associated with chronic gut-derived peripheral inflammation (Fig. 4 H, I).

Concluding, the integration of multi-omics data revealed a strong correlation between Factor 1, which is significantly elevated in chronic DSS colitis mice, and midbrain microglia. These findings strengthen the above reported findings that microglia may critically contribute to disease pathology and could represent a promising therapeutic target.

### Chronic colitis persists despite delayed Csf1r inhibition

We next applied an interventional approach to delineate the role of microglia in orchestrating the midbrain immune response and potential neurotoxicity in chronic colitis. As all microglial cluster showed high expression levels of *Csf1r* with only a slight decrease in Cluster 3 (Fig. 4C), we fed mice with chow supplemented with the Csf1r antagonist PLX5622 to efficiently reduce microglial numbers. To ensure the onset of colitis prior to treatment, PLX5622 administration was started on day 11, after the first of three DSS cycles (Fig. 5A). PLX5622-fed mice developed colitis and extraintestinal inflammation of similar severity compared to mice receiving placebo chow (Fig. 5B-I). Histologic assessment of hematoxylin and eosin-stained colon showed excessive immune cell infiltration into the mucosa and loss of epithelial barrier integrity, resulting in mucosal thickening and adding up to a higher histologic score in chronic DSS colitis mice independent of Csf1r inhibition (Fig. 5F). Colonic Iba1^+^ myeloid cells showed a trend toward increased abundance, although this change did not reach statistical significance due to substantial interindividual variability. Co-administration of PLX5622 failed to significantly reduce Iba1⁺ cell density (Fig. 5I). These data indicate that targeting Csf1r-dependent myeloid cells by PLX5622 treatment after the first DSS treatment cycle does not influence major hallmarks of chronic DSS colitis.

**Figure 5.**
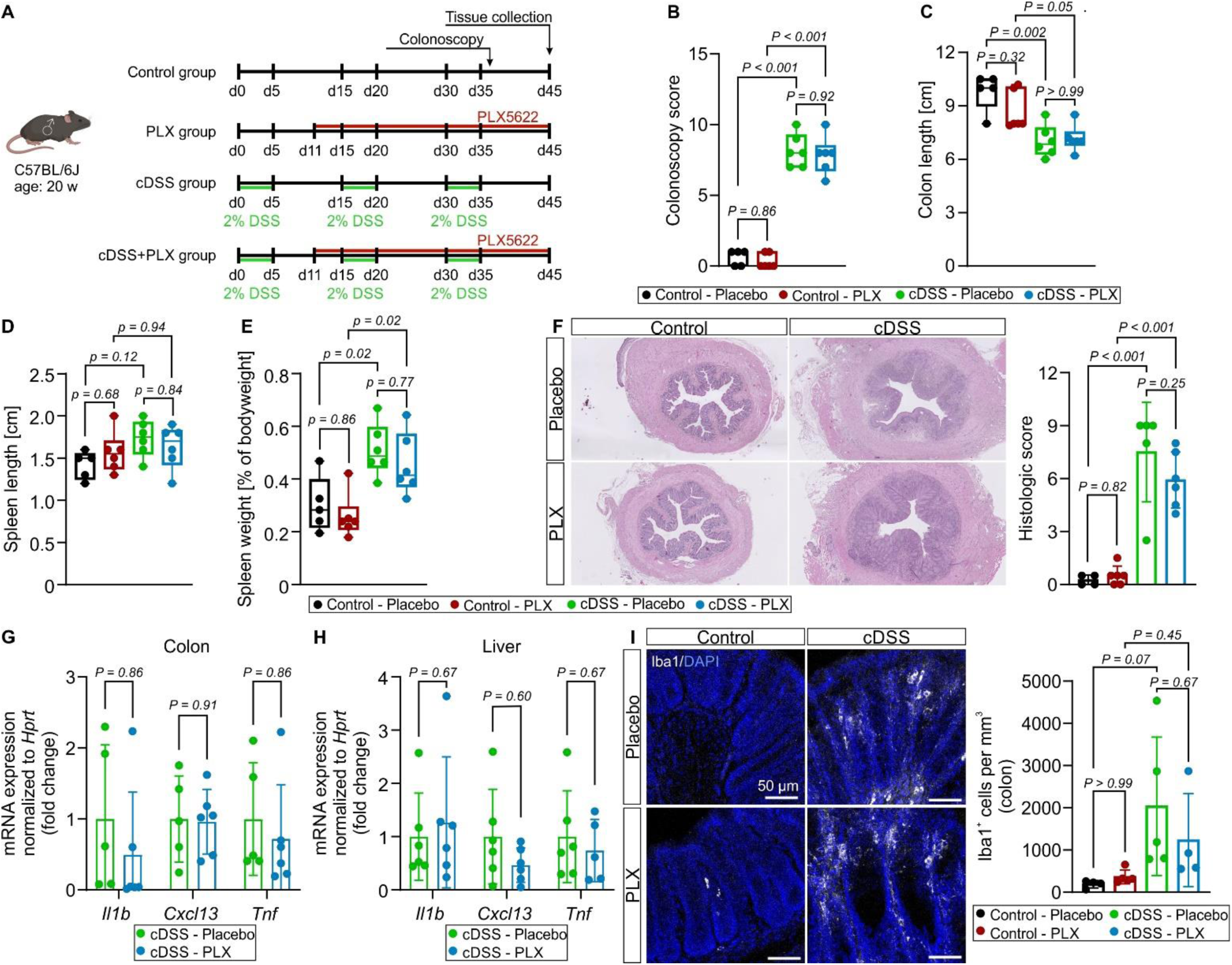
Csf1r-targeting of myeloid cells after colitis onset does not alter chronic colitis and systemic inflammation progression. **A** Timeline of colitis induction and PLX5622 administration in male C57BL/6J mice aged 20 weeks at the start of the experiment. **B** Colonoscopy score on day 36. **C** Colon length measured post dissection. **D** Spleen length measured post dissection. **E** Spleen weight in percentage of bodyweight. **F** Representative images of H&E-stained colon sections (left) and histologic score (right). **G**, **H** Gene expression analysis of the inflammation-associated genes *Il1b*, *Cxcl13*, and *Tnf* in the colon (**G**) and liver (**H**) (multiple unpaired *t*-tests). **I** Immunostaining for Iba1 (white, left) and quantification of Iba1^+^ cells normalized to mm^3^ in the colon (right). All data represent *n* = 4-6 mice per group. Two-way ANOVA with Tukey’s multiple comparisons test, if not otherwise indicated. Data are presented as mean ± s.d. Each dot represents the value of an individual mouse. Icons in **A** were created with BioRender.com.

### Csf1r-dependent microglial depletion rescues dopaminergic neuron loss

Next, we investigated the impact of PLX5622 treatment on the CNS of mice with chronic DSS colitis and controls. We performed flow cytometry of whole brain tissue and specifically investigated the midbrain using immunofluorescence (Fig. 6A). PLX5622 treatment for 35 days led to a profound reduction of CD45^int^CD11b^+^ cells in the whole brain by 63% or 34% when comparing PLX5622- to placebo-fed control or chronic DSS colitis mice, respectively (Fig. 6B). Focusing on the midbrain, we detected a significant reduction in Iba1^+^ microglia numbers by 67% in PLX5622-vs. placebo-fed chronic DSS colitis mice (Fig. 6C). Assessing the impact of myeloid cell targeting on the mixed midbrain immune cell infiltration, we found unchanged levels of CD3⁺ T cells and CD8⁺ T cells between PLX5622- and placebo-fed chronic DSS colitis mice by flow cytometry and immunofluorescence (Fig. 6D, E). Also, the percentage of Ly6G^+^ neutrophils of CD45^high^ cells in whole brain tissue and the number of vessel-associated Ly6G^+^ cells in the midbrain were comparable between chronic DSS colitis mice receiving placebo or PLX5622-supplemented chow (Fig. 6F, G).

**Figure 6.**
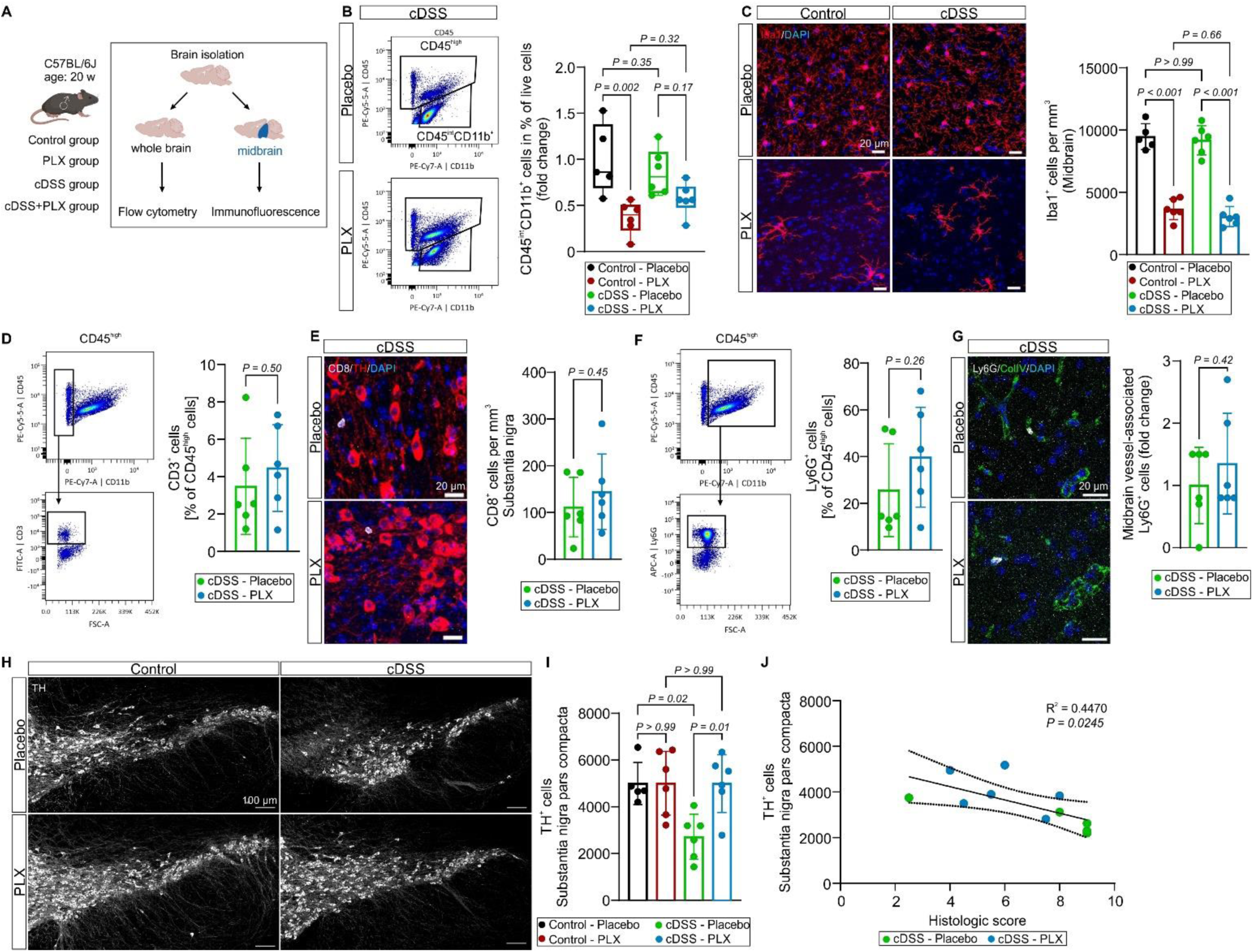
Myeloid cell targeting after colitis onset efficiently reduces microglia in the midbrain and rescues dopaminergic neuron loss in the substantia nigra pars compacta. **A** Schematic representation of whole brain and midbrain analysis using flow cytometry and immunofluorescence, respectively. **B** Gating of CD45^int^CD11b^+^ cells (left) and percentage of CD45^int^CD11b^+^ cells of live cells (right) analyzed by flow cytometry. **C** Immunostaining for Iba1 (red, left) and quantification of Iba1^+^ cells in the midbrain normalized to mm^3^ (right). **D** Flow cytometry analysis of CD3^+^ immune cells in whole brain tissue (left) and proportion of CD3^+^ cells of CD45^high^ cells (right). **E** Immunostaining for CD8 (white) and tyrosine hydroxylase (TH, red) in chronic DSS colitis mice (left) and quantification of CD8^+^ cells in the substantia nigra normalized to mm^3^ (right; two-tailed, unpaired *t*-test). **F** Flow cytometry analysis of Ly6G^+^ immune cells in whole brain tissue (left) and proportion of Ly6G^+^ cells of CD45^high^ cells (right). **G** Immunostaining for Ly6G (white) and ColIV (green, left) and quantification of vessel-associated Ly6G^+^ cells in the midbrain (right; two-tailed, unpaired *t*-test). **H** Immunostaining for TH (white). **I** Quantification of all TH^+^ cells in the total substantia nigra pars compacta of one brain hemisphere. **J** Correlation and linear regression analysis of the TH^+^ cell number in the substantia nigra pars compacta (**I**) and the histologic score (Fig. 4F) in PLX5622- and placebo-fed chronic DSS colitis mice. The results **B** - **J** are from *n* = 5-6 mice per group. Two-way ANOVA with Tukey’s multiple comparisons test, if not otherwise indicated. Data are presented as mean ± s.d. Each dot represents the value of an individual mouse. Icons in **A** were created with BioRender.com.

Finally, we wondered if dopaminergic neuron loss in the substantia nigra of adult mice with chronic colitis can be alleviated by Csf1r-dependent microglial depletion. Intriguingly, PLX5622 treatment during the second and third cycle of chronic DSS colitis led to a complete rescue of TH-expressing neurons in the substantia nigra pars compacta (Fig. 6H, J). The numbers of dopaminergic neurons correlated with the severity of colitis (Fig. 6J).

In summary, Csf1r inhibition after colitis onset in the chronic DSS colitis model results in a significant reduction of microglia in the midbrain and prevents an inflammation-mediated loss of dopaminergic neurons in the substantia nigra pars compacta without affecting colitis severity and migration of bone marrow-derived immune cells to the midbrain.

## Discussion

Our study fundamentally advances the understanding of brain disease in the context of gut inflammation. We show a causal link between chronic colitis-induced neuroinflammation and the loss of dopaminergic neurons. Despite the complex neuroinflammatory fingerprint in the midbrain, this neurodegenerative process is predominantly driven by microglia and profoundly rescued by Csf1r inhibition – independent of colitis severity. The potential impact of chronic gut inflammation on the development of neuroinflammation and neurodegenerative diseases is particularly relevant in PD, given its association with IBD and early, prominent gastrointestinal symptoms^[6, 37, 38]^. To better model the association of chronic colitis with later-onset neurodegeneration, we chose a chronic paradigm in mice at an age of 20 weeks for adult-onset colitis induction, in contrast to most previous studies investigating DSS colitis in 2-3-month-old mice^[10, 39]^. Moreover, we aimed to model a chronic gut inflammation by the relapsing remitting DSS treatment paradigm over three cycles. Our approach led to a 46% loss of dopaminergic neurons in the substantia nigra pars compacta, while previous studies with shorter DSS treatment paradigms reported either a less pronounced loss of approximately 35%^[11]^ or no reduction of dopaminergic neurons by DSS treatment alone^[39]^. The loss of dopaminergic neurons in the present model is also supported by the downregulation of GO terms related to mitochondrial function in midbrain proteomic data of mice with chronic DSS colitis, as a similar signature was recently identified in the substantia nigra of PD patients by spatial transcriptomics^[40]^.

As indicated by previous studies, there is mounting evidence of a regional brain immune response to gut inflammation^[12]^, as well as a central role for inflammatory microglia in neurodegeneration^[33]^. Here we performed an in-depth characterization of the neuroimmune response paired with multi-omics analyses and revealed its causal relation to neuropathology in the context of chronic gut inflammation. We identified a susceptibility of the midbrain to chronic gut inflammation on gene and protein level and determined a complex innate and adaptive immune cell response in the midbrain using single-cell RNA-sequencing in the chronic DSS colitis mouse model of IBD.

The regional responsiveness of midbrain microglia described here is in line with a study conducted two decades ago reporting an increased activation of microglia in the midbrain upon local administration of lipopolysaccharide (LPS), which was accompanied by a loss of dopaminergic neurons, whereas cortical and hippocampal microglia were less reactive to local LPS^[41]^. Furthermore, the midbrain was reported to be a preferential entry site for blood-derived immune cells in the context of peripheral inflammation and experimental autoimmune encephalomyelitis^[42]^.

Our single-cell profiling of the midbrain immune response to chronic gut inflammation revealed a shift from homeostatic to inflammatory microglia including an expansion of an interferon-response cluster (Cluster 10) as well as an increased migration of bone marrow-derived immune cells into the midbrain parenchyma and adjacent compartments. Of note, CD8^+^ T cells expressing *Nkg7* were linked to neurodegeneration in the context of Alzheimer’s Disease (AD)^[43]^. Also in PD, CD8^+^ T cells are present in the substantia nigra early in disease and may co-contribute to dopaminergic neuron loss^[44, 45]^. In addition, neutrophil adhesion to brain capillaries was previously implicated in memory loss via regulation of cerebral blood flow in a mouse model of AD^[46]^. However, our integrative analysis points to a predominant role of microglia in the midbrain response to chronic colitis. Indeed, by our interventional approach, we confirm that microglia are essential for dopaminergic neuron loss in chronic gut inflammation, as microglial reduction by 67% using PLX5622 treatment protected dopaminergic neurons without affecting the migratory capacity of CD8^+^ T cells or neutrophils towards the midbrain and adjacent blood vessels. Apart from direct neurotoxic effects, it is conceivable that microglia signal to CD8^+^ T cells or neutrophils via direct interaction or secreted molecules, as microglial interaction with T cells and neutrophils were recently implicated with neurodegeneration in AD and tauopathy, respectively^[47, 48]^. Within the microglial compartment, interferon-response microglia are highly conserved in different models of neurodegeneration^[33]^ and may specifically contribute to dopaminergic neuron loss, as microglial type-I interferon signaling was recently detected on single-cell RNA-sequencing datasets of human PD-patient-derived midbrain samples^[49]^.

Our data further imply that myeloid cell targeting by Csf1r inhibition directly acts in the CNS to protect dopaminergic midbrain neurons. PLX5622 treatment did not affect endoscopic and structural hallmarks of colitis. In line with this, the oral CSF1R inhibitor PRV-6527 failed to meet clinical endpoints in a Phase 2a clinical trial in patients with Crohn’s Disease^[50]^. While we found no significant reduction of total Iba1^+^ myeloid cells in the colon upon PLX5622 treatment, previous data on the impact of Csf1r inhibition on gut immune cells are heterogenous^[51–53]^. In contrast, midbrain microglia were strongly reduced upon PLX5622 treatment, which was linked to a rescue of dopaminergic neurons. Thus, Csf1r inhibition may have no substantial impact on gut pathology in IBD, but may effectively protect from IBD-related neurologic comorbidity, particularly PD in later life, by mitigating microglial numbers. Furthermore, our treatment approach with administration of PLX5622 during the second and third DSS treatment cycle suggests that the first cycle of DSS treatment is not sufficient to induce dopaminergic neuron loss, indicating that rather chronic sustained gut-derived inflammation is required to elicit neurodegenerative processes in the midbrain.

Collectively, we identified microglia as a potential target to rescue dopaminergic neuron loss in the context of IBD. Our findings are in line with previous epidemiological observations in humans revealing an increased risk for patients with IBD to develop PD, while chronic anti-inflammatory treatment with TNF inhibitors restored the risk in IBD patients^[38]^. Future studies comparing midbrain immune signatures of IBD patients and PD patients, in particular those suffering from the recently proposed body-first PD subtype^[54, 55]^, will reveal the importance of our findings in human disease by shared and disease-specific mechanisms along the gut-immune-brain axis.

## Acknowledgements

We thank the Core Unit für Zellsortierung und Immunomonitoring, Friedrich-Alexander-Universität Erlangen-Nürnberg, Erlangen, Germany for performing single cell sorting. We thank Julia Derdau for preparation of single-cell RNA-sequencing libraries. Access to the MESO QuickPlex SQ 120MM instrument was kindly provided by the Translational Radiobiology (Department of Radiation Oncology, Universitätsklinikum Erlangen).

## Author information

Authors and Affiliations

**Department of Molecular Neurology, Universitätsklinikum Erlangen, Friedrich-Alexander-Universität Erlangen-Nürnberg, Erlangen, Germany**

Rebecca Katharina Kutscherauer, Wie Xiang, Alexander Grotemeyer, Jürgen Winkler, Patrick Süß

**Department of Medicine 1, Universitätsklinikum Erlangen, Friedrich-Alexander-Universität Erlangen-Nürnberg, Erlangen, Germany**

Iris Stolzer, Emely Elisa Neumaier, Mark Dedden, Sebastian Zundler, Claudia Günther

**Department of Neurology, Universitätsklinikum Erlangen, Friedrich-Alexander-Universität Erlangen-Nürnberg, Erlangen, Germany**

Veit Rothhammer

**Department of Chemistry, Ludwig-Maximilians-Universität München, München, Germany**

Pavel Kielkowski

**Institute of Neuropathology, Medical Faculty, University of Freiburg, Freiburg, Germany**

Marco Prinz, Klaus-Peter Knobeloch

**Signalling Research Centres BIOSS and CIBSS, University of Freiburg, Freiburg, Germany**

Marco Prinz, Klaus-Peter Knobeloch

**Division of Molecular Neuroimmunology, Medical Institute of Bioregulation, Kyushu University, Fukuoka, Japan**

Takahiro Masuda

**Department of Neurosciences, University of California, San Diego, USA**

Johannes Schlachetzki

**Deutsches Zentrum Immuntherapie, Universitätsklinikum Erlangen, Erlangen, Germany**

Veit Rothhammer, Sebastian Zundler, Jürgen Winkler, Claudia Günther, Patrick Süß

## Contributions

R.K.K. performed and analyzed most of the experiments and wrote the manuscript. R.K.K., S.Z., W.X., J.S., J.W., C.G., and P.S. conceived the project and designed experiments. I.S., E.E.N., A.G., and P.S. contributed to mouse experiments. J.S. performed bulk RNA-sequencing. R.K.K. and J.S. analyzed bulk RNA-sequencing data. P.K. performed mass spectrometry. R.K.K. and P.K. analyzed proteomic data. R.K.K. and M.D. analyzed single-cell RNA-sequencing data. M.P., T.M., and K.-P.K. provided *Hexb^tdTomato^* mice. C.G. performed histologic scoring of colon sections. J.W., C.G., and P.S. supervised the research. P.S. acquired data and wrote the manuscript. M.P., T.M., V.R., S.Z., and J.W. provided advice and interpreted the data. J.W., C.G., and P.S. contributed to directing and funding of the study. R.K.K. prepared the figures. V.R., S.Z., and J.W. substantially revised the manuscript.

## Corresponding author

Patrick Süß

