## Supplementary Figures for "Csf1r-mediated depletion of midbrain microglia prevents dopaminergic neuron loss during chronic colitis"

Rebecca Katharina Kutscherauer et al.

**Supplementary Figures**

**Figure S1.**

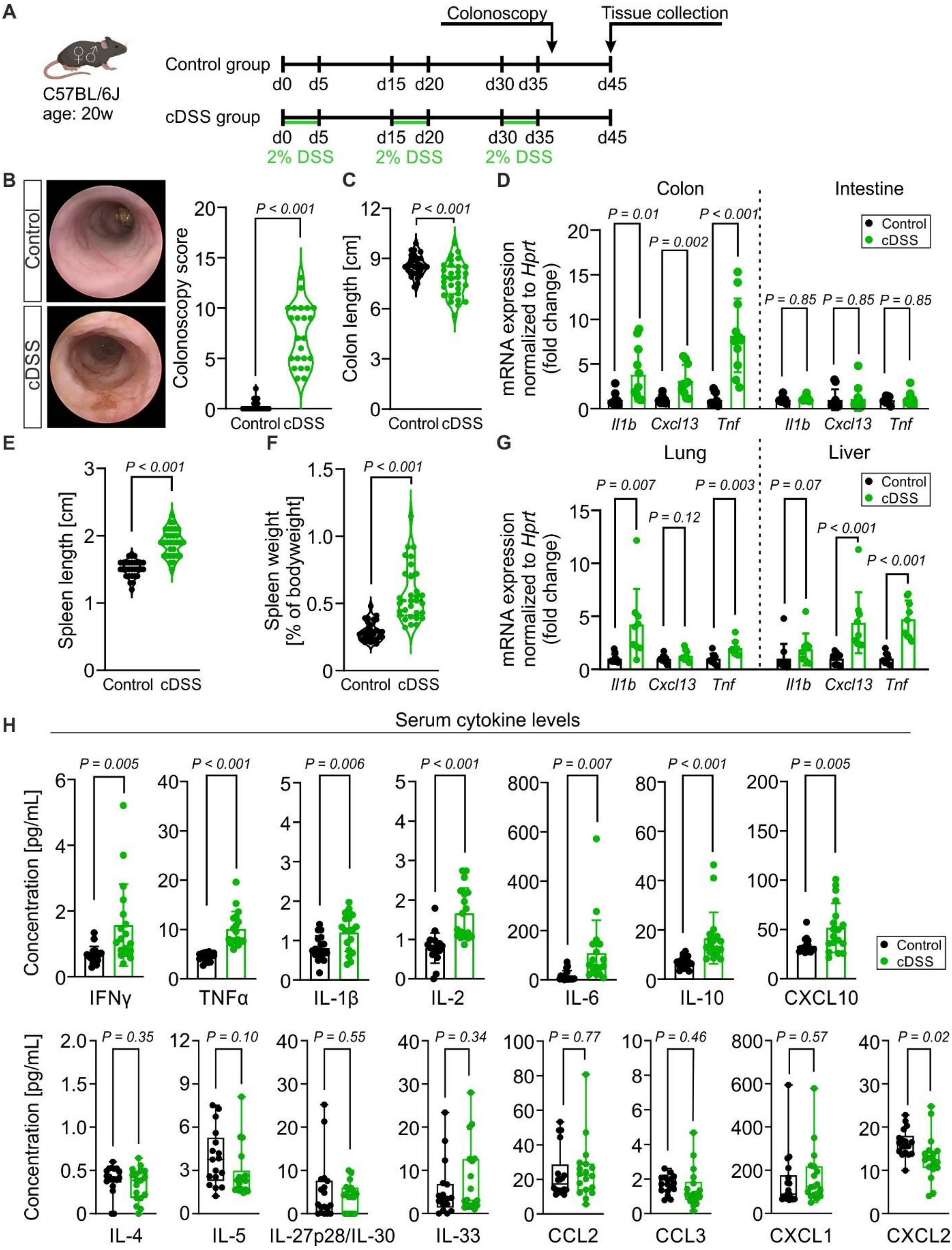


**Figure S1. Cyclic DSS treatment induces chronic colitis and systemic inflammation in C57BL/6J mice.**

**A** Timeline of colitis induction in male and female C57BL/6J mice aged 20 weeks at the start of the experiment. **B** Representative colonoscopy images and colonoscopy scores (*n* = 23 mice per group; pooled of three independent experiments). **C** Colon length post dissection (*n* = 29-30 mice per group; pooled of four independent experiments). **D** Expression of the inflammation-associated genes *Il1b*, *Cxcl13*, and *Tnf* in the colon and small intestine (*n* = 10-11 mice per group; multiple unpaired *t*-tests). **E**, **F** Spleen length and spleen weight in percentage of bodyweight (*n* = 29-31 mice per group; pool of four independent experiments). **G** Expression of the inflammation-associated genes *Il1b*, *Cxcl13*, and *Tnf* in the lung and liver (*n* = 9-10 mice per group; multiple unpaired *t*-tests). **H** Serum concentrations of the inflammation-associated cytokines IFNγ, TNFα, IL-1β, IL-2, IL-6, IL-10, CXCL10, IL-4, IL-5, IL-27p28/IL-30, IL-33, CCL2, CCL3, CXCL1 and CXCL2 (*n* = 18 mice per group). Two-tailed, unpaired *t*-test, if not otherwise indicated. Each point represents the value of one mouse. Data are presented as mean ± s.d. Icons in **A** were created with BioRender.com.

**Figure S2.**

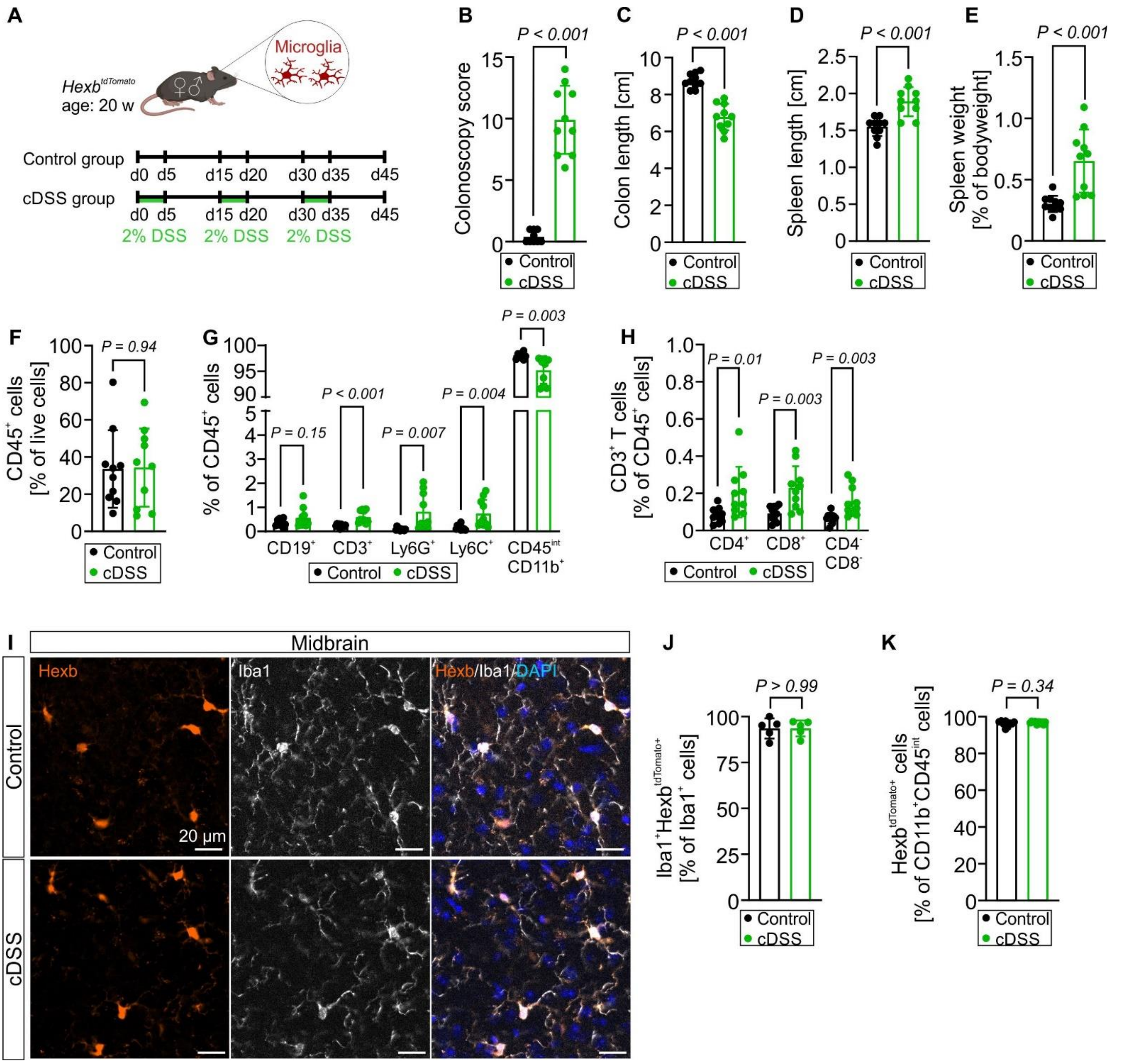


**Figure S2. Cyclic DSS treatment induces chronic colitis and immune cell response in the whole brain of *Hexb*$^{tdTomato}$ mice while microglial Hexb$^{tdTomato}$ expression is maintained. A** Timeline of colitis induction in male and female *Hexb*$^{tdTomato}$ mice aged 20 weeks at the start of the experiment. **B** Colonoscopy score on day 36. **C** Colon length post dissection. **D** Spleen length post dissection. **E** Spleen weight in percentage of bodyweight. **F**, **G**, **H** Flow cytometry analysis of CD45$^+$ immune cells (**F**), CD19$^+$ B cells, CD3$^+$ T cells, Ly6G$^+$ neutrophils, Ly6C$^+$ monocytes, and CD45$^{int}$CD11b$^+$ microglia as percentage of $CD45^+$ cells (**G**), and $CD3^+CD4^+$, $CD3^+CD8^+$, and $CD3^+CD4^-CD8^-$ T cells as percentage of $CD45^+$ cells (**H**) in whole brain tissue. **I**, Immunostaining for Iba1 (white) and Hexb (tdTomato) in brain tissue of *Hexb*$^{tdTomato}$ mice. Scale bar, 20 µm. **J**, Quantification of $Iba1^+Hexb^{tdTomato+}$ cells in percentage of $Iba1^+$ cells in the midbrain. **K**, Flow cytometry analysis of $Hexb^{tdTomato+}$ cells in percentage of $CD11b^+CD45^{int}$ cells. The results are from $n$ = 10 (**B - H**, **K**) and $n$ = 5 (**J**) mice per group. Two-tailed, unpaired $t$-test, if not otherwise indicated. Each point represents the value of one mouse. Data are presented as mean ± s.d. Icons in **A** were created with BioRender.com.

**Figure S3.**

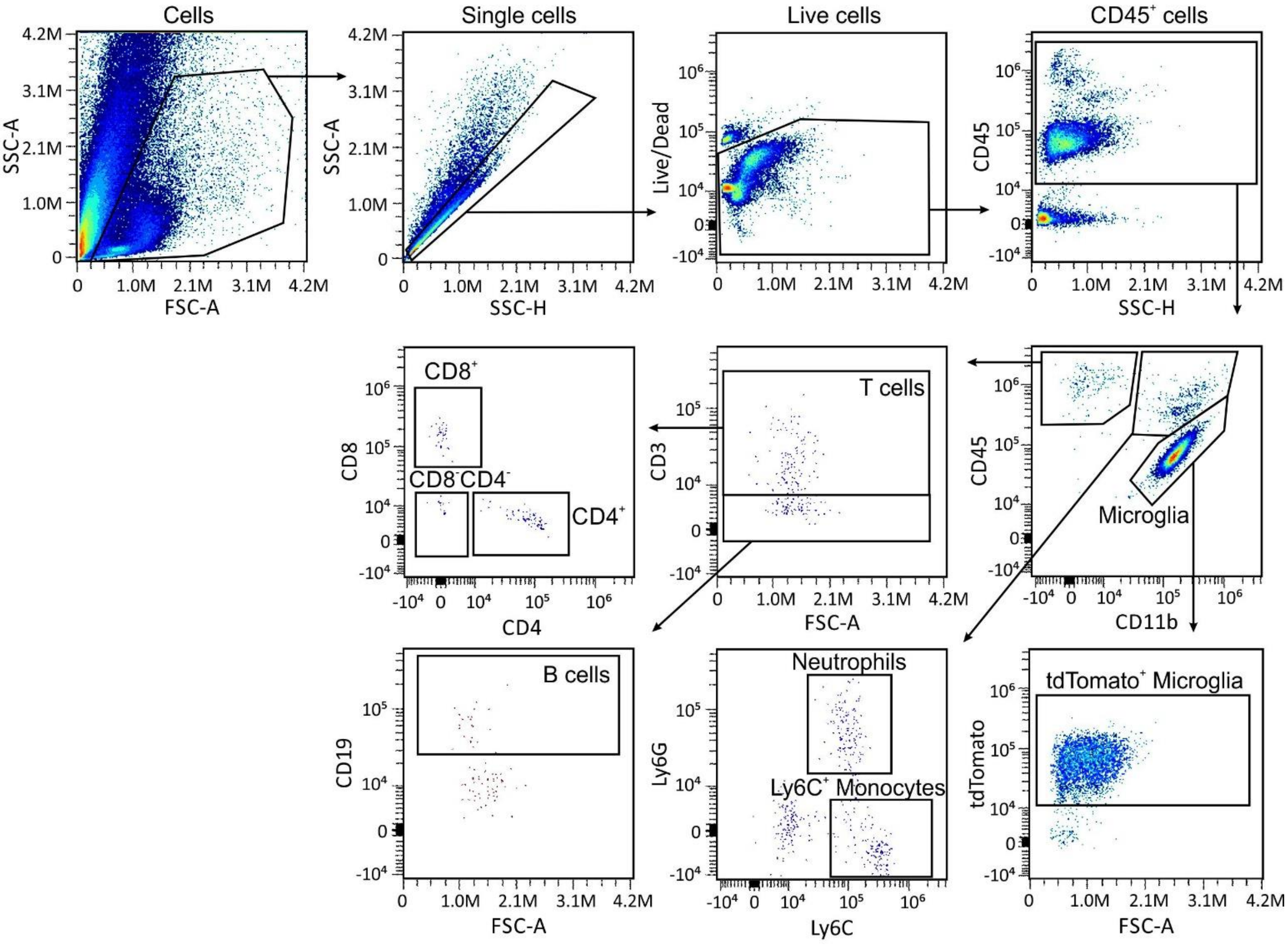


**Figure S3. Flow cytometry gating strategy of brain immune cells isolated from cDSS-treated and control *Hexb*$^{tdTomato}$ mice.** Cells were gated on SSC-A and SSC-H to remove doublets. Singlets negative for the Live/Dead fixable Aqua stain were considered as live cells. $CD45^+$ live cells were gated on CD11b to identify $CD11b^+CD45^{int}$ microglia that were further gated on tdTomato to determine $tdTomato^+$ microglia. $CD11b^+CD45^+$ cells were gated on Ly6G and Ly6C to distinguish $Ly6G^+$ neutrophils from $Ly6C^+$ monocytes. $CD11b^-CD45^+$ cells were gated on CD3 (T cells) and CD19 (B cells), and $CD3^+$ cells were further gated on CD8 and CD4 to identify T cell subpopulations.

**Figure S4.**

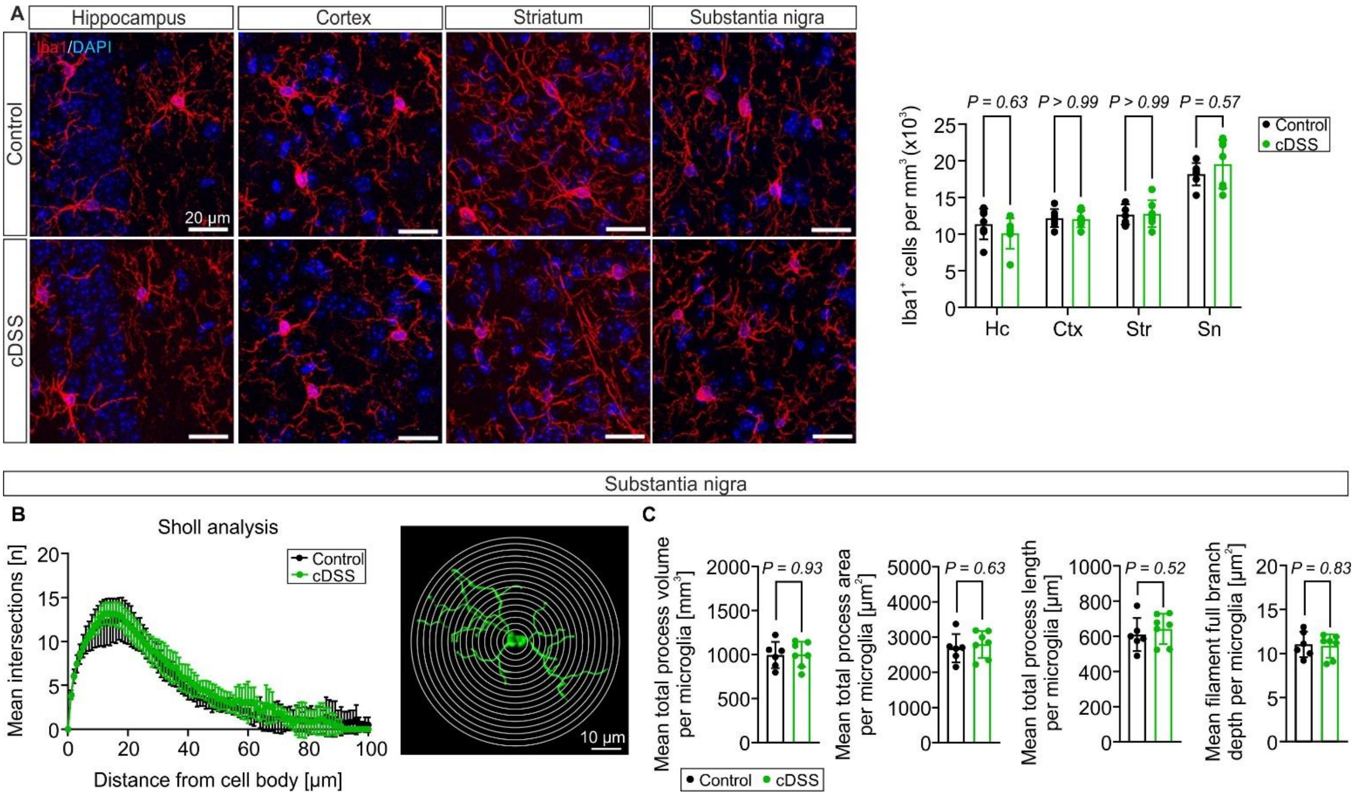


**Figure S4. Regional microglia density and morphology remain unaltered in chronic DSS colitis mice.**

**A** Immunostaining for Iba1 in brain tissue of C57BL/6J mice (left) and quantification of Iba1$^+$ cells normalized to mm$^3$ in the hippocampus (Hc), cortex (Ctx), striatum (str), and substantia nigra (Sn, $n$ = 7 mice per group, two-way ANOVA with Šídák's multiple comparisons test). Scale bars, 20 µm. **B** Sholl analysis of microglia in the substantia nigra. **C** Microglial process parameters in the substantia nigra ($n$ = 6-7 mice per group; two-tailed, unpaired $t$-test). Each point represents the mean value of one mouse. Data are presented as mean ± s.d.

**Figure S5.**

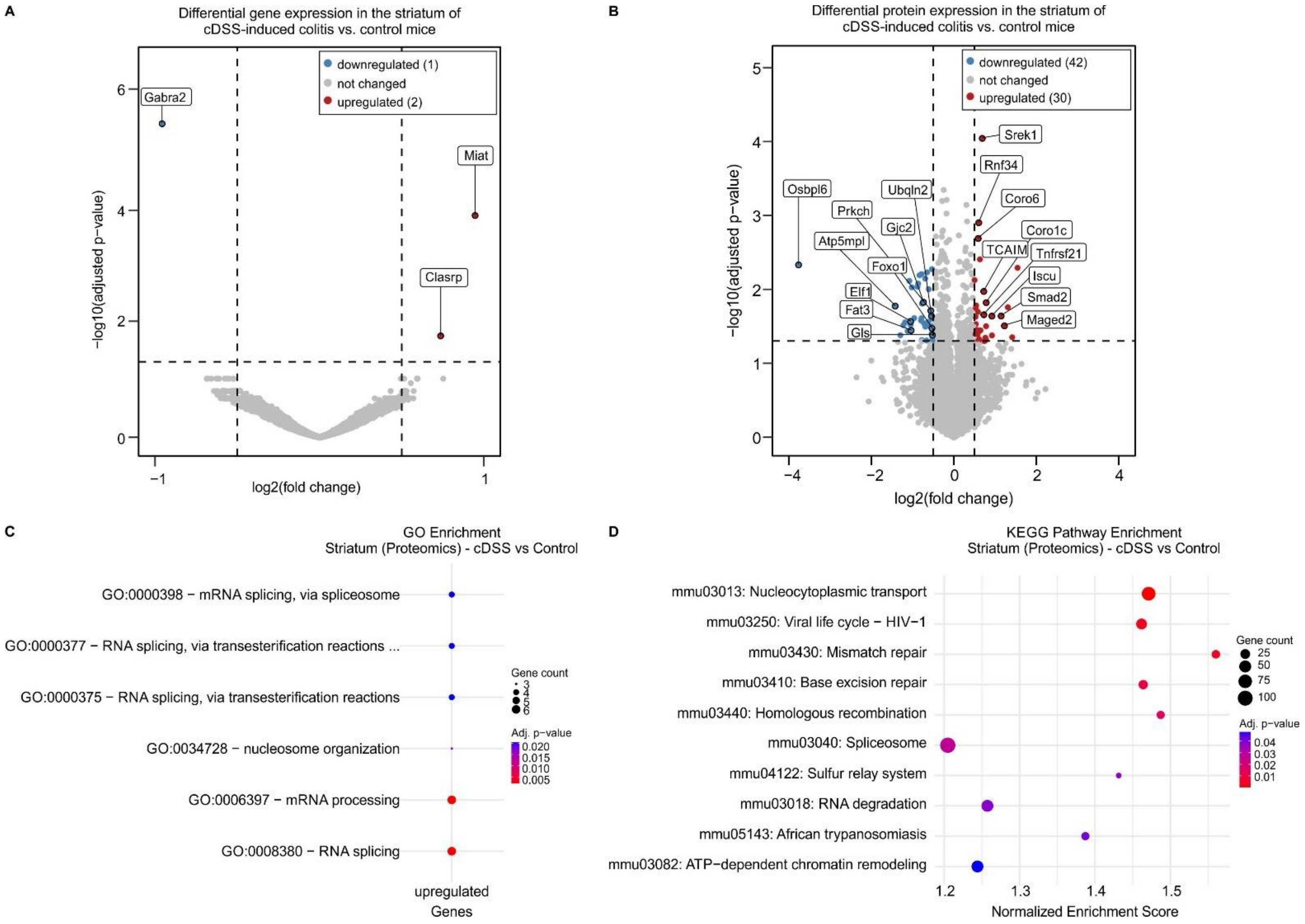


**Figure S5. Transcriptome and proteome analyses reveal subtle alterations in the striatum during chronic DSS colitis. A** Volcano plot of genes differentially expressed in the striatum of chronic DSS colitis mice vs. control mice. **B** Volcano plot of proteins differentially expressed in the striatum of chronic DSS colitis mice vs. control mice. **C** Gene Ontology Biological Process analysis of the 30 proteins significantly higher expressed in the striatum of chronic DSS colitis mice compared to control mice. **D** KEGG (GSEA) pathway enrichment of proteins differentially expressed in the striatum of chronic DSS colitis mice compared to control mice.

**Figure S6.**

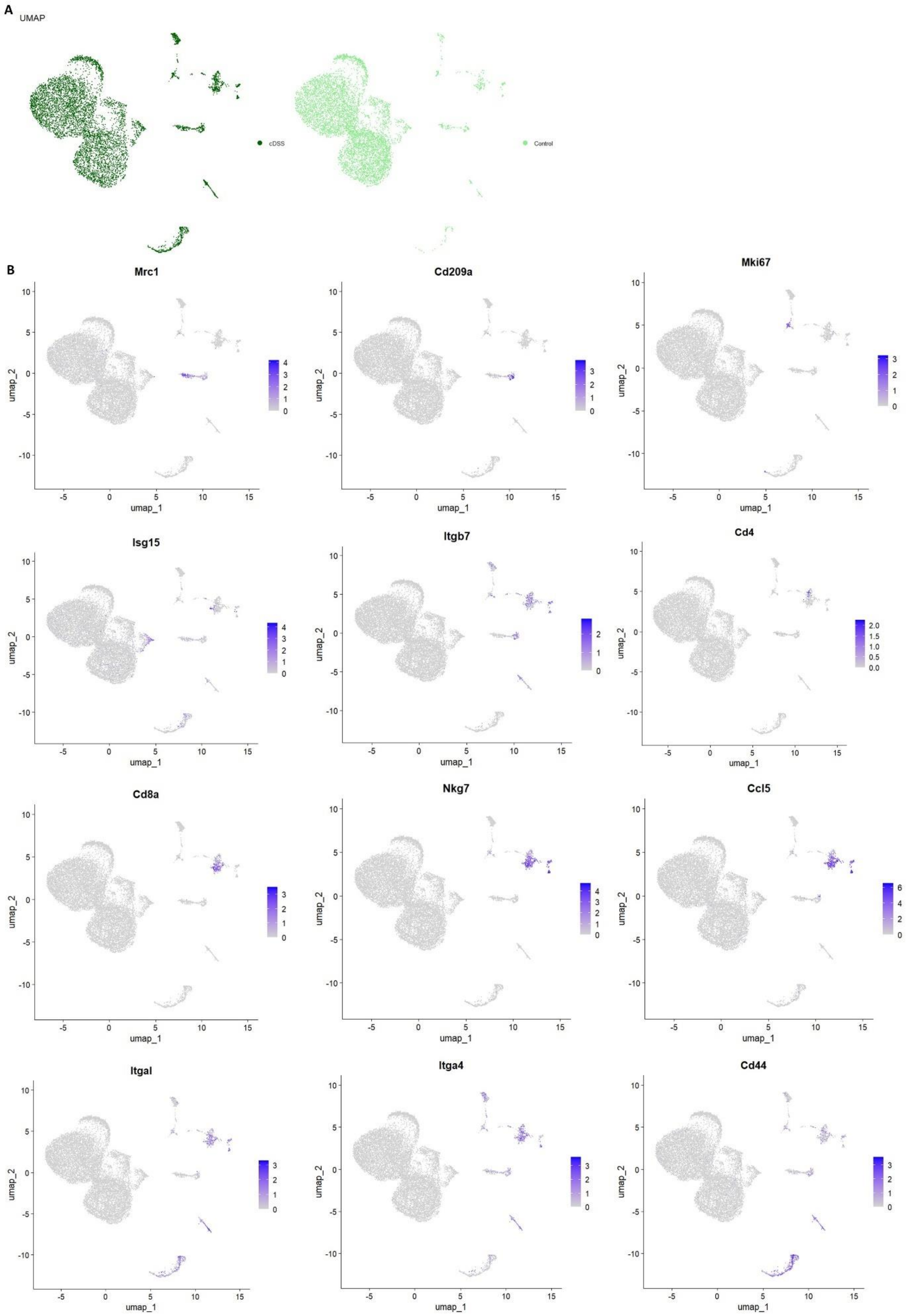
A
UMAP
cDSS
Control
B
Mrc1
Cd209a
Mki67
Isg15
Itgb7
Cd4
Cd8a
Nkg7
Ccl5
Itgal
Itga4
Cd44
umap_1
umap_2

**Figure S6. UMAPs and feature plots of immune cell type defining genes in the midbrain. A** Individual UMAPs of immune cells in the midbrain per group. **B** Feature plots of cluster defining markers.

80
