## Supplementary Table for "Csf1r-mediated depletion of midbrain microglia prevents dopaminergic neuron loss during chronic colitis"

Rebecca Katharina Kutscherauer et al.

**Supplementary Tables**

**Table S1. Primer sequences for reverse transcription-quantitative PCR.**

| Gene | Primer forward (5' - 3') | Primer reverse (5' - 3') |
|---|---|---|
| *Cxcl13* | CTCCAGGCCACGGTATTCTG | CCAGGGGGCGTAACTTGAAT |
| *Hprt* | GTCATGTCGACCCTCAGTCC | GCAAGTCTTTCAGTCCTGTCC |
| *Il1b* | GCAACTGTTCCTGAACTCAACT | ATCTTTTGGGGTCCGTCAACT |
| *Pgk1* | GTCGTGATGAGGGTGGACTT | AACGGACTTGGCTCCATTGT |
| *Tnf* | TAGCCCACGTCGTAGCAAAC | GCAGCCTTGTCCCTTGAAGA |
